# In vivo genome-wide CRISPR screens in human T cells to enhance T cell therapy for solid tumors

**DOI:** 10.1101/2025.09.23.678127

**Authors:** Qi Liu, Peixin Amy Chen, Esha Urs, Shimin Zhang, Maya M. Arce, Charlotte H. Wang, Zhongmei Li, Jin Seo, Nupura Kale, Taylor N. LaFlam, Fanglue Peng, Eric Shifrut, Greg M. Allen, Justin Eyquem, Katherine Fuh, Stacie E. Dodgson, Jason G. Cyster, Alexander Marson, Julia Carnevale

## Abstract

Large-scale CRISPR screening in human T cells holds significant promise for identifying genetic modifications that can enhance cellular immunotherapy. However, many genetic regulators of T cell performance in solid tumors may not be readily revealed in vitro. In vivo screening in tumor-bearing mice offers greater physiological relevance, but has historically been limited by low intratumoral T cell recovery. Here, we developed a new model system that achieves significantly higher human T cell recovery from tumors, enabling genome-wide in vivo screens with small numbers of mice. Tumor-infiltrating T cells in this model exhibit hallmarks of dysfunction compared to matched splenic T cells, creating an ideal context for screening for genetic modifiers of T cell activity in the tumor microenvironment. Using this platform, we performed two genome-wide CRISPR knockout screens to identify genes regulating T cell intratumoral abundance and effector function (e.g., IFN-γ production). The intratumoral abundance screen uncovered the P2RY8-Gα13 GPCR signaling pathway as a negative regulator of human T cell infiltration into tumors. The effector function screen identified GNAS (Gαs), a central signaling mediator downstream of multiple GPCRs that sense different suppressive ligands, as a key regulator of T cell dysfunction in tumors. Targeted GNAS knockout rendered T cells resistant to multiple suppressive cues and significantly improved therapeutic performance across diverse solid tumor models. Moreover, combinatorial knockout of P2RY8 (trafficking) and GNAS (effector function) further enhanced overall tumor control, demonstrating that genetic modifications targeting distinct T cell phenotypes can be combined to improve therapeutic potency. This flexible and scalable in vivo screening platform can be adapted to diverse tumor models and pooled CRISPR libraries, enabling future discovery of genetic strategies that equip T cell therapies to overcome barriers imposed by solid tumors.

## Main

T cell-based immunotherapies have revolutionized the treatment of hematologic malignancies, with remarkable clinical success in leukemias and lymphomas. However, despite recent regulatory approvals in select solid tumors, the broader application of T cell therapies to the majority of solid tumors, which account for most cancer cases, remains a major unmet challenge^1,2^. Solid tumors pose a unique set of obstacles to T cell therapies, such as restricted T cell infiltration, metabolic constraints, immune suppression within the tumor microenvironment (TME), and more^3,4^. Overcoming these barriers requires innovative strategies to engineer T cells capable of navigating the hostile conditions in solid tumors. Recent advances in CRISPR-based approaches offer unprecedented opportunities to identify and manipulate gene targets that can reprogram T cells to resist or bypass these challenges^5–8^. Despite this potential, most genetic screens have been confined to in vitro systems, which cannot replicate all of the complex and hostile conditions of the TME. For instance, in vitro models fail to recapitulate the nutrient and oxygen deprivation encountered by T cells in vivo, and they do not adequately capture the complex immunosuppressive milieu, which includes inhibitory cytokines and chemokines, suppressive metabolites, suppressive cells, and immune checkpoint ligands that can substantially impair T cell efficacy.

Taking a reductionist approach to modeling the T cell’s experience in tumors, a number of groups have attempted to design in vitro studies to simulate various elements found in the TME for CRISPR screening. For instance, we previously performed a variety of different in vitro genome-wide human T cell screens in parallel, adding various suppressive factors individually, including suppressive cytokines, suppressive metabolites, small molecules to mimic suppressed states, and suppressive cell types (Tregs) to identify gene knockout targets that confer resistance to each of these suppressive factors^6^. We used a cross-comparative analysis to determine those gene targets that engender resistance to specific screen conditions, and those that appeared to confer resistance across multiple suppressive factors. Other groups have performed in vitro CRISPR screens in human T cells where repeated co-culture with tumor cells is used to model the T cell dysfunction that arises in the setting of chronic antigen exposure^9,10^. Additionally, CARs with high tonic signaling have been used to model T cell exhaustion pressures in in vitro screens^7^.

There have been early efforts to conduct pooled genetic screens in human T cells within tumor-bearing immunodeficient mice, but these have generally been limited in scale due to the low numbers of T cells recoverable from tumors. In a prior collaborative study, for example, we screened a focused library of 48 sgRNAs targeting 20 genes in vivo^11^. The library size was constrained by the limited recovery of NY-ESO-1 TCR transduced human T cells from A375 melanoma tumors. Typically, only ∼50,000 to 500,000 human T cells can be isolated per tumor, which poses a substantial barrier to genome-wide screening. Standard libraries contain ∼75,000 sgRNAs (e.g., 4 sgRNAs per gene), and achieving 1000x coverage would require approximately 300 NSG mice per donor, amounting to ∼600 mice per screen across two donors, rendering such experiments logistically impractical. Similar limitations have been encountered in murine T cell screens using immunocompetent tumor-bearing models, where only ∼25,000 OT-I T cells can typically be recovered per tumor^12,13^. As a result, achieving high-coverage screens (>500x) with large-scale libraries of tumor-bearing mice, while maintaining robust sgRNA representation, has remained a major technical hurdle.

To overcome this limitation, we developed a novel in vivo CRISPR screening platform that enables genome-scale loss-of-function screens in primary human T cells within tumor-bearing mice. This system, which involves overexpression a CD3 scFv on tumor cells, allows high recovery of tumor-infiltrating T cells, permitting comprehensive analysis with a relatively small number of animals. Using this approach, we conducted the first genome-wide in vivo CRISPR screens in human T cells, identifying gene targets that regulate both T cell abundance and effector function in solid tumors. Notably, these screens revealed significant enrichment for genes involved in GPCR signaling. We focused on two GPCR pathways that regulate (1) T cell trafficking and infiltration, and (2) T cell dysfunction in response to suppressive ligands within the tumor microenvironment. Genetic disruption of these pathways enhanced both intratumoral accumulation and resistance to immunosuppressive cues. Moreover, these targets can be edited individually or in combination to significantly improve the efficacy of T cell therapies across multiple solid tumor models. Together, these results demonstrate that our in vivo screening model enables discovery of therapeutically relevant biology that would likely be missed in conventional in vitro systems.

## Results

### Development of an in vivo model system enabling large-scale human T cell screens in a small number of tumor-bearing mice

To improve T cell recovery from tumors in mice, we developed a new system where we engineered A375 melanoma cells to express components of an anti-CD3 scFv **(Fig. 1a and Extended Data Fig. 1a)**, such that the tumor cells can engage and activate any polyclonal repertoire of CD8 and CD4 T cells. We isolated 70 single-cell clones following lentiviral transduction of A375 melanoma cells with the anti-CD3 scFv construct, allowing us to avoid polyclonal heterogeneity and instead select clones with uniform CD3 scFv expression for consistent T cell activation in downstream assays (**Extended Data Fig. 1b-c**). We subsequently narrowed the selection to 27 clones based on anti-CD3 scFv expression levels, where clones were selected to cover the full range of anti-CD3 scFv expression levels observed (**Extended Data Fig. 1d**). We evaluated the ability of primary human T cells to kill these 27 single tumor cell clones in vitro and identified three clones with varying degrees of susceptibility to killing by human T cells (**Extended Data Fig. 1e,** labeled low, medium, and high). While these three clones we selected for study exhibited similar growth rates to wild-type (WT) A375 cells, we noted that many of the selected clones showed slower tumor growth than WT A375 cells and these were not selected for further study. We engrafted the three selected A375 clones individually into the flanks of NSG mice and treated 11 days later with polyclonal human T cells **(Extended Data Fig. 1f)**. These clones with different anti-CD3 scFv densities generated a range of responses to human T cells in vivo, parallelling the range of killing susceptibilities we observed in vitro **(Extended Data Fig. 1g)**.

**Fig. 1.**
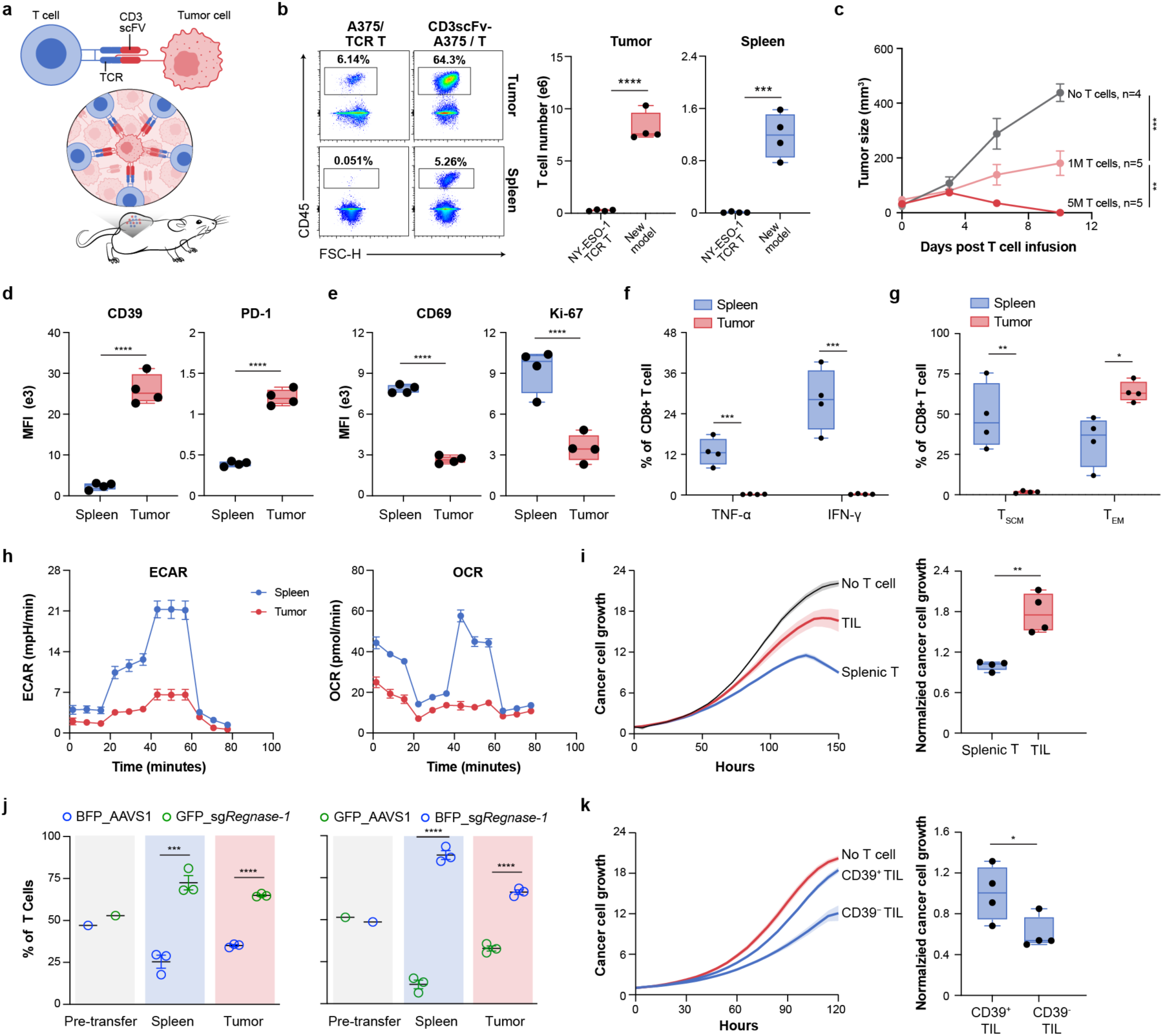
Anti-CD3 scFv-expressing tumor model enables efficient recovery of human T cells from tumors and induces hallmarks of intratumoral T cell dysfunction. a. Schematic of the in vivo model utilizing anti-CD3 scFv-overexpressing tumors for T cell enrichment in solid tumors. b. Flow cytometry gates (left) and absolute counts (right) of NY-ESO-1 TCR transduced T cells isolated from A375 wild-type tumors and spleens versus polyclonal T cells from A375^low^ tumors expressing anti-CD3 scFv and spleens in NSG mice. n = 4 mice for each group. c. Average tumor growth curves of A375^low^ tumor-bearing NSG mice treated with T cells 11 days after tumor engraftment. n = 4 mice for no T cell control, 5 mice for both 1M T cell and 5M T cell groups. **d-g.** Comparative analysis of T cell phenotypes in spleens versus tumors 9 days post-treatment in A375^low^ tumor-bearing NSG mice (n = 4 mice for each group). Shown are: mean fluorescence intensity (MFI) of CD39 and PD1 **(d)** and CD69 and Ki-67 (**e**) in CD8⁺ T cells; percentage of TNF-α⁺ and IFN-γ⁺ CD8⁺ T cells **(f)** and percentage of stem cell memory (Tscm) and effector memory (Tem) CD8⁺ T cells (**g**). **h-i.** Functional and metabolic profiling of splenic T cells versus TILs. Seahorse analysis of extracellular acidification rate (ECAR) and oxygen consumption rate (OCR) (**h**, representative data from one of three mice), and Incucyte-based killing assay (**i**, n = 4 mice per group) are shown. **j.** In vivo competition assay of AAVS1 control and Regnase-1 knockout primary human T cells. Primary human T cells transduced with either a control AAVS1 sgRNA or an sgRNA targeting Regnase-1 were mixed at a 1:1 ratio and co-transferred into A375^low^ tumor-bearing mice. The relative proportions of each T cell population in the spleen and tumor were analyzed 14 days after infusion. Left: BFP_AAVS1 control sgRNA mixed with GFP_Regnase-1 sgRNA; Right: GFP_AAVS1 control sgRNA mixed with BFP_Regnase-1 sgRNA. n = 3 mice for each group. **k.** Incucyte-based killing assay comparing sorted CD39^+^ versus CD39^-^ TILs isolated from A375^low^ tumors and then co-cultured with A375^low^ tumor cells ex vivo (n = 4 mice). *P* values were determined by two-tailed unpaired Student’s t-test (**b, d-g, i-k**) and two-way ANOVA (**c**) \**P* < 0.05, \*\**P* < 0.01, \*\*\**P* < 0.001, and \*\*\*\**P* < 0.0001. Data are presented as mean ± s.e.m.

To preserve a dynamic range in which we can observe improvements in tumor cell killing, we subcutaneously implanted the A375^low^ and A375^medium^ anti-CD3 scFv expressing A375 tumor cells into mice and then injected these mice with polyclonal T cells from healthy human donors. After 9 days, we isolated T cells from tumors and spleens. We routinely recovered approximately 7 million T cells per tumor in the A375^low^ line and around 30 million T cells per tumor in the A375^medium^ line. In contrast, only approximately 200 thousand T cells were recovered from each subcutaneous WT A375 melanoma tumor in NSG mice treated with engineered NY-ESO-1– specific human TCR T cells (**Fig. 1b and Extended Data** Fig 1h). Importantly, T cells isolated from A375^low^ tumors demonstrated higher levels of canonical exhaustion markers (CD39), and lower levels of activation (CD69) compared to those from A375^medium^ tumors (**Extended Data Fig. 1i-j**). We elected to move forward with the A375^low^ cell line for further experiments because it struck the ideal balance between achieving high T cell recovery while promoting T cell dysfunction, thus better modeling exhausted tumor infiltrating lymphocytes (TILs). Despite evidence of dysfunction, transferred donor T cells can slow or eradicate these A375^low^ tumors at standard (1M) and high (5M) doses, respectively (**Fig. 1c**), further suggesting that this model strikes a balance between activating and challenging the tumor-infiltrating T cells. As expected for a robust TIL model, the A375^low^ tumor-derived T cells also show evidence of exhaustion and dysfunction when compared to the splenic T cells isolated from the same mice. In addition to expressing higher levels of canonical surface markers of exhaustion (PD-1, CD39), the TIL expressed lower levels of activation markers (CD154, CD69, CD137) and were less proliferative (Ki-67) compared to splenic T cells (**Fig. 1d**-**e** **and Extended Data Fig. 2a-b**). The TIL also demonstrated lower effector cytokine production, lower stem cell-like memory T cell states, and impaired metabolic function (assayed by Seahorse and flow metrics) compared to splenic T cells (**Fig. 1f-h and Extended Data Fig. 2c-f**). Finally, TIL isolated from tumors exhibited reduced ex vivo tumor cell killing compared to splenic T cells (**Fig. 1i**). Thus, in this model, TIL experience conditions that induce dysfunction similar to the T cell states observed in patient TIL^14^, suggesting that this in vivo system could be used to screen for genes that mitigate such dysfunctional states.

To evaluate the ability of this model system to distinguish gene modifications that enhance T cell competitive fitness in vivo, we performed a pooled competition assay comparing T cells edited for a positive control gene (ZC3H12A/Regnase-1 knockout) to those edited at a negative control safe harbor locus (AAVS1). Regnase-1 has been shown to be a potent negative regulator of T cell expansion and persistence in tumors and spleens in vivo^12^. To test whether our new model system could recapitulate these findings, we performed a competition assay using human T cells transduced with AAVS1 control or ZC3H12A-targeting sgRNA vectors encoding distinct fluorescent reporters, followed by nucleofection with Cas9 protein. We transferred a 50:50 mixture of BFP_AAVS1: GFP_Regnase-1 KO or GFP_AAVS1: BFP_Regnase-1 KO T cells into our tumor-bearing mice and isolated T cells from spleens and tumors 14 days later (**Extended Data** Fig 2g). As expected, Regnase1-KO T cells showed a robust competitive advantage across T cell donors in both tumors and spleens, suggesting our model system can be used to distinguish gene targets known to boost T cell expansion and persistence in vivo (**Fig. 1j, Extended Data Fig. 2h-i**).

In addition to using sgRNA abundance to measure the fitness and persistence of gene edits in vivo, we sought to enable functional screening by flow sorting tumor-infiltrating T cells based on phenotypic markers. To this end, we isolated T cells from tumors and sorted them based on expression of CD39, a well-established marker of exhaustion resulting from chronic antigen stimulation^15^. We found that compared to the CD39-high sorted TIL from this model, the CD39-low TIL showed superior tumor killing in a tumor co-culture assay ex vivo (**Fig. 1k, Extended Data Fig. 2j)**. These findings demonstrate that this platform not only enables robust recovery of TILs from tumors, but also allows for functional stratification of screened populations by flow cytometry to identify sgRNAs that drive distinct T cell states.

### In vivo genome-wide CRISPR loss-of-function screen identifies GPCR signaling regulators of T cell enrichment in solid tumors

Using this novel in vivo screening platform, we conducted a genome-wide CRISPR knockout (KO) screen in human T cells from two donors, quantifying sgRNA abundance in T cells recovered from tumors and spleens relative to their representation at the time of injection. T cells from two donors were isolated from PBMCs, activated with CD3/CD28 beads, transduced with lentivirus to deliver the Brunello genome-wide (GW) library of sgRNAs, and expanded in culture for 11 days (**Fig. 2a**). This library contains approximately 75,000 sgRNAs, which requires recovery of roughly 38-75 million T cells to achieve 500-1000x coverage. NSG mice were subcutaneously injected with 2 million A375^low^ tumor cells, and 17 days later, each mouse was injected with 4 million pooled edited T cells from one of the two donors. Pooled edited T cells were injected across 11 mice for each donor (22 mice total), achieving ∼600x library coverage per donor at the time of T cell injection. After 10 days, tumors and spleens were harvested, and 80-100 million T cells were isolated for each human T cell donor (>1000x coverage) and pooled for gDNA isolation and next generation sequencing. The 10-day collection time-point was selected based on the Regnase-1 KO competition experiment, where enrichment became detectable after day 9 (**Extended Data Fig. 3a**). We assessed relative sgRNA frequencies in T cells recovered from the tumors and spleens of mice compared to the T cell population at the time of injection. Analysis of overall sgRNA coverage from T cells isolated from tumors and spleens for each donor showed the uniform distributions in each condition (GINI index scores ranged from 0.0542-0.069, **Extended Data Fig. 3b**). In addition, strong correlation was observed for sgRNAs from the tumors and spleens compared to input for each donor, suggesting high coverage and no library bottleneck events (**Extended Data Fig. 3c**).

**Fig. 2.**
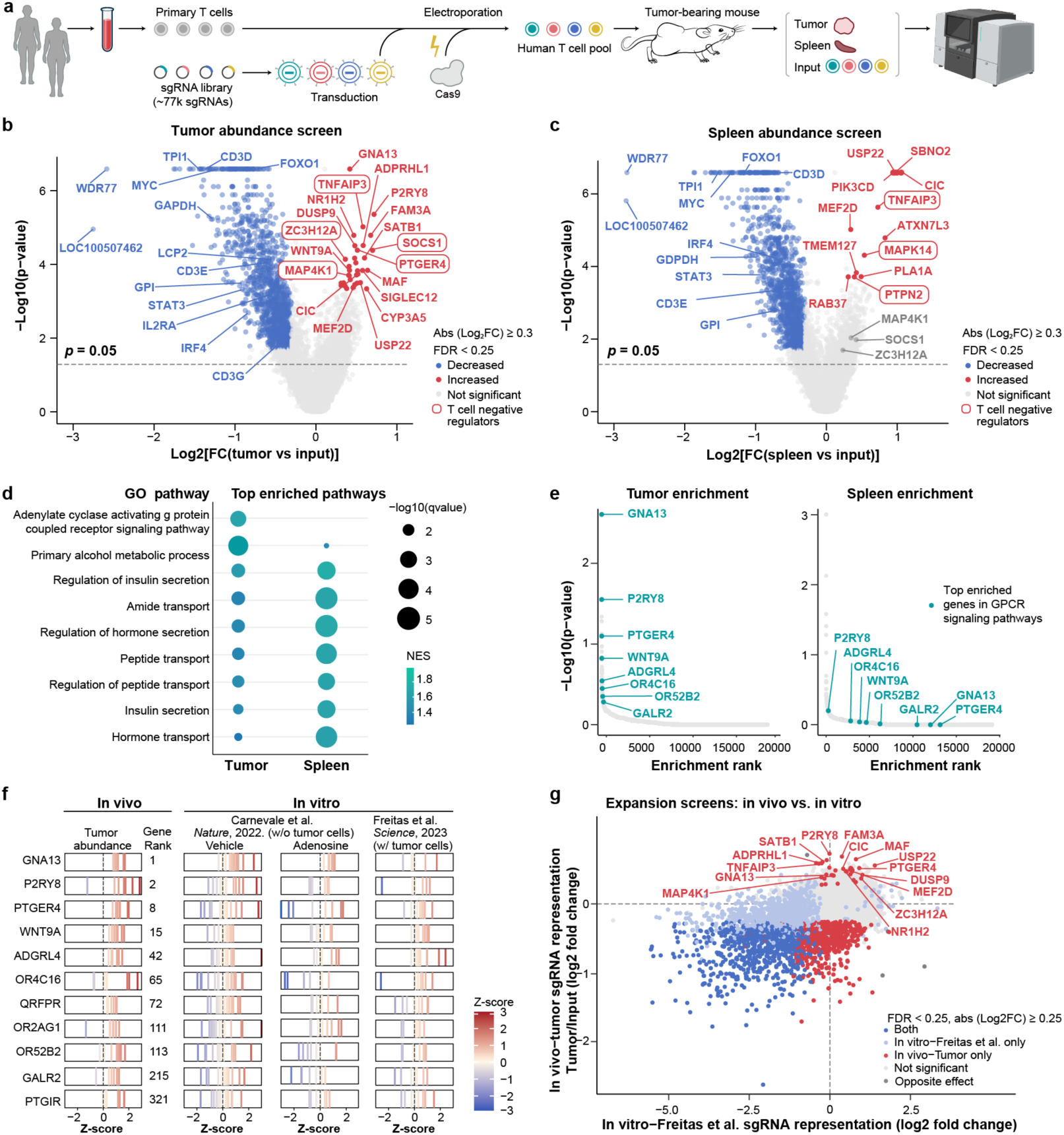
Genome-wide in vivo CRISPR screen identifies G protein-coupled receptor signaling pathways regulating human T cell abundance in solid tumors. **a.** Schematic of the in vivo genome-wide CRISPR screen strategy for regulators of T cell abundance in tumors. **b-c.** Volcano plots indicate genes with enriched (red) and depleted (blue) guides in T cells from tumors (**b**) and spleens (**c**) identified by MAGeCK for tumor vs input and spleen vs input. Each screen includes two human T cell donors. (Enrichment based on FDR < 0.25, abs (Log2FC)≥0.3). Selected enriched genes with known roles in human T cell fitness are circled. Top enriched GO biological processes pathways in the tumor vs input and spleen vs input comparisons, based on top normalized enrichment score (NES) and cutoff of min(qvalue) < 0.0005. Gene rank plots of in vivo tumor (tumor versus input) and spleen (spleen versus input) abundance CRISPR screens. Colored points highlight the top 250 genes from the tumor abundance screen that are associated with the GPCR signaling pathways. sgRNA z-scores for genes of interest that are associated with the GPCR signaling pathways, shown across the in vivo tumor abundance screen and published in vitro screens^6,7^. Comparison of gene-level log_2_ fold changes from genome-wide CRISPR knockout screens in human T cells. Each point represents a gene. Log_2_ fold change from the in vivo T cell abundance screen (y-axis) is plotted against log_2_ fold change from a published in vitro expansion screen (x-axis; Freitas et al.⁷).

MAGeCK analysis-derived rank scores for sgRNAs enriched in the tumors compared to input showed enrichment for many gene targets with previously well-defined roles in enhancing human T cell antitumor immunity upon knockout (**Fig. 2b, Extended Data Table 1**). For instance, ZC3H12A (Regnase-1), SOCS1, PTGER4, MAP4K (HPK1) and TNFAIP3 have all been published as bona fide targets to enhance human T cell antitumor efficacy^12,16–19^. sgRNAs targeting these established target genes were also enriched to lower extents in the spleen abundance screen, suggesting that either these genes are enhancing systemic proliferation or that there is some transit from the tumor to the spleen (**Fig. 2c**, labeled in grey). Depletion of sgRNAs targeting genes with known positive roles in T cell activation and function such as LCP2, STAT3, IL2RA, and components of the CD3 receptor was also evident in the tumor and spleen datasets, (negative LFC, **Fig. 2b-c**). KEGG and Gene Ontology (GO) pathway analyses also confirmed dropout in pathways essential for T cell viability and function (**Extended Data Fig. 3d-e**). Furthermore, a number of high-ranking genes with sgRNAs enriched in tumors have known roles in mouse T cell differentiation, dysfunction, and exhaustion, such as TNFAIP3, SATB1, MEF2D, PTGER4 and MAF (**Fig. 2b)**^20–25^. Thus, our screen successfully identified established T cell regulators based on differential abundance in vivo.

To gain insights into functional pathways enriched in our screens, we completed gene set enrichment analysis (GSEA) to identify GO biological processes pathways enriched among the tumor and spleen abundance screen hits. These analyses showed specific enrichment for G-Protein-Coupled Receptor (GPCR) signaling pathway mediators in hits from the T cells isolated from tumors compared to spleens, while a variety of metabolic pathways were found to be enriched in both screens (**Fig. 2d and Extended Data Fig. 3f**). Because genes from the GPCR signaling pathway gene set were enriched among top-ranking genes in the tumor but not in the splenic samples, the data suggest a unique role for GPCR activity in T cells within the tumor (**Fig. 2e**). Comparative sgRNA level analysis of enriched GPCR pathway genes between our in vivo dataset and previously published in vitro screens designed to identify regulators of T cell dysfunction^6,7^ demonstrated more uniformly positive enrichment in our in vivo screen for GPCR components (**Fig. 2f**), some of which have already been shown to play roles in TIL dysfunction^18,21,26^. To evaluate the specificity of our in vivo T cell abundance screen more comprehensively, we compared our tumor enrichment data with results from a previously published in vitro screen of human T cell expansion using a tonically signaling GD2 CAR. In that study, T cells underwent tumor cell challenges in culture to drive expansion under conditions of chronic stimulation^7^. This comparison highlighted unique enrichment of sgRNAs for various genes in our in vivo tumor condition that were not enriched in the in vitro screen conditions using a common cutoff of FDR < 0.25 (**Fig. 2g**). These genes span a range of biological processes, such as MAP kinase signaling (e.g. DUSP9, MAP4K1), GPCR signaling (e.g. PTGER4, P2RY8, GNA13), transcriptional regulation (e.g. MEF2D, CIC, USP22, MAF, SATB1), deubiquitinase activity (e.g. TNFAIP3, USP22), T cell lineage commitment (e.g. SATB1, MAF), and lymphocyte trafficking (e.g. P2RY8, GNA13, SATB1). Overall, this comparison suggests that performing screens for T cell abundance in vivo can highlight gene targets with important roles for T cell accumulation and persistence in tumors that may be missed by in vitro screens.

### The P2RY8 signaling pathway inhibits human T cell trafficking in solid tumors

We compared gene target enrichment in the tumor to the spleen to highlight gene KOs that were uniquely enriched in the tumor (**Fig. 3a**). Of the top 50 ranked genes based on tumor versus spleen enrichment, 12% have known roles in GPCR signaling or previously defined roles in cell trafficking and migration (S1PR1, GNA13, CXCR4, ARHGEF1, PTGER4, and P2RY8)^27–33^ (**Extended Fig 4a**). Of these, we noted that P2RY8, GNA13 (Gα13), and ARHGEF1 are all components of the P2RY8 GPCR signaling pathway, a key pathway known to regulate B cell migration in the germinal centers of lymphoid organs^29,30^ (**Fig. 3b**). P2RY8 signaling inhibits migration of human germinal center B cells and T follicular helper cells towards chemoattractants and it is thought to be critical in promoting cell confinement in germinal centers^34^. While P2RY8 signaling has been primarily studied in regulating B cell migration, it has also been shown that this pathway can negatively regulate B and T cell migration into the bone marrow^35^. Our screen results suggest that this pathway may also be relevant in T cell trafficking in cancer. Importantly, while P2RY8 is conserved across many vertebrate species, it lacks a functional orthologue in mice^30^ and therefore would not have been identified in a fully murine in vivo screen (**Fig. 3c**). This underscores a key advantage of our platform: the ability to uncover functional roles for human-specific genes in human T cells within the context of tumor-bearing mice.

**Fig. 3.**
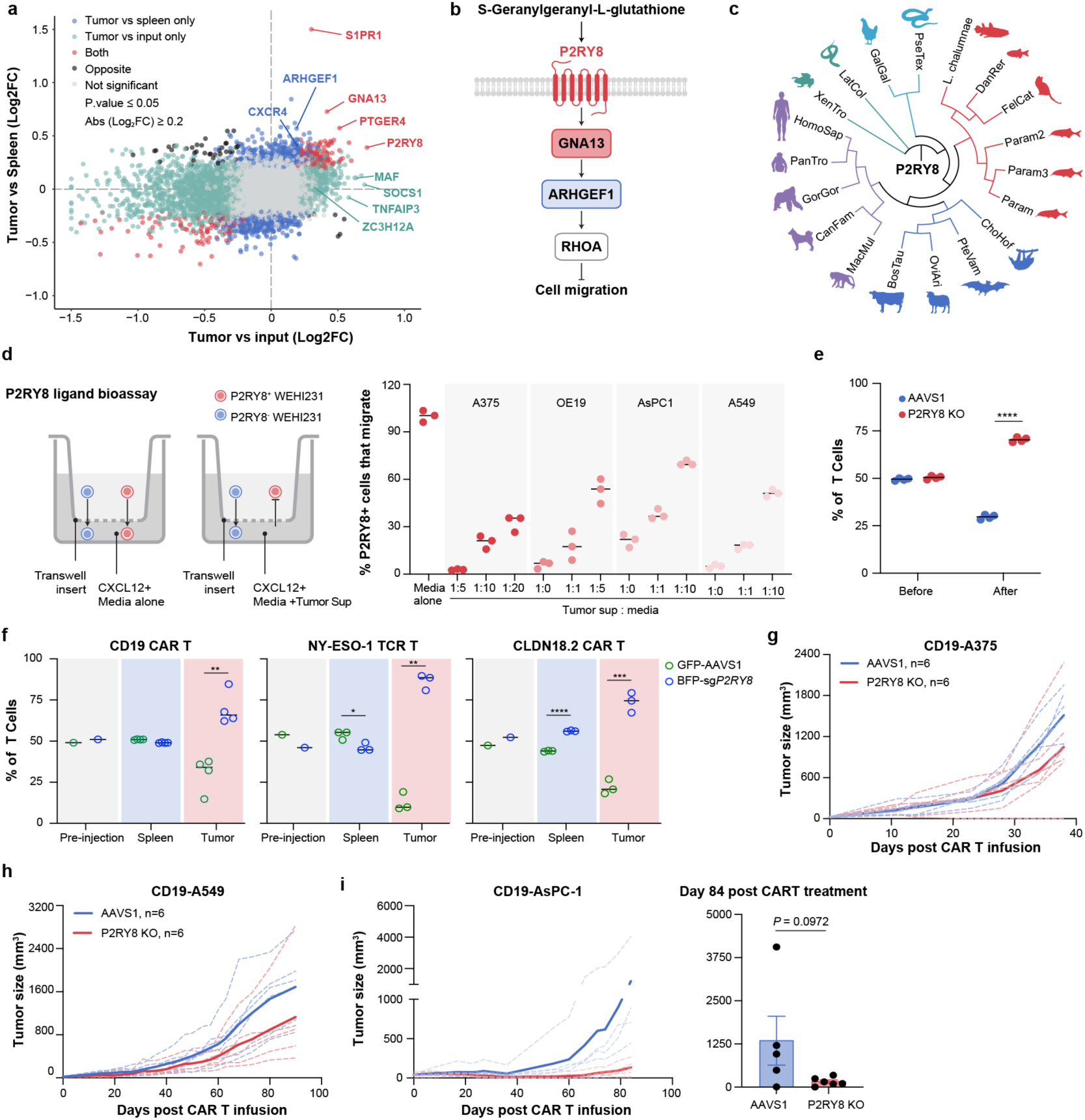
GGG-P2RY8 axis inhibits human T cell trafficking in solid tumors. Comparison of gene enrichment values (log_2_ fold change, across two donors) in the in vivo abundance screen, evaluating tumor versus spleen and tumor versus input conditions, colored by direction and specificity of effect comparing the two conditions. Genes meeting the significance threshold (p-value<=0.05 and abs (Log2FC)≥0.2) are highlighted. a. Schematic of the P2RY8 signaling pathway that inhibits germinal center B cell migration. Schematic is based on literature^34^. b. Phylogenetic tree of P2RY8 homologues across different species, with rodents notably absent. c. In vitro P2RY8 ligand bioassay. Schematic (left): GFP-P2RY8-expressing and wild-type WEHI-231 cells were placed in the upper chamber of a transwell insert. Migration toward CXCL12 in the lower chamber was quantified by flow cytometry, measuring the percentage of GFP-P2RY8-expressing cells that reached the bottom well. Representative data (right): Migration is normalized to GFP-P2RY8-expressing cells that migrated toward migration media alone. Tumor supernatants were diluted with fresh migration media at the indicated ratios (n = 3 biological replicates per group). d. In vitro human T cell transwell migration assay. AAVS1 control and P2RY8 KO human T cells were labeled with CellTrace™ Violet and CellTrace™ Far Red, respectively, and mixed at a 1:1 ratio. Flow cytometry analysis showing the percentages of AAVS1 and P2RY8 KO human T cells within total CD45⁺ T cells before being added to the transwell insert (before) and after migration to the bottom well (after). Data represent four technical replicates and are representative of one of three independent human T cell donors. e. In vivo migration assay. GFP-labeled AAVS1 control and BFP-labeled P2RY8 KO CD19 CAR T cells, NY-ESO-1 TCR T cells, or CLDN18.2 CAR T cells were mixed at a 1:1 ratio and intravenously co-transferred into NSG mice bearing CD19-A375, A375, or OE19 tumors, respectively. Forty-eight hours after transfer, the distribution of GFP⁺ AAVS1 and BFP⁺ P2RY8 KO T cells in spleen and tumor were analyzed by flow cytometry (n = 3-4 mice per group). **g-h.** Tumor growth in NSG mice bearing CD19-A375 tumors (**g**) or CD19-A549 tumors **(h)** treated with AAVS1 vs P2RY8 KO CD19 CAR T cells 9 days post tumor injection (n = 6 mice for each group). **i.** Tumor growth of CD19-AsPC-1 tumor-bearing NSG mice treated with AAVS1 vs P2RY8 KO CD19 CAR T cells 25 days post tumor injection (left) and tumor size at day 84 post CAR T cell infusion (right) (n = 5 mice for AAVS1 control and 6 mice for P2RY8 KO). *P* values were determined by two-tailed unpaired Student’s t-test (**e, f, i**). \**P* < 0.05, \*\**P* < 0.01, \*\*\**P* < 0.001, and \*\*\*\**P* < 0.0001. Data are presented as mean ± s.e.m.

Our screen data suggest that P2RY8 signaling may inhibit T cell migration into tumors. It has been shown that the metabolite *S*-Geranylgeranyl-L-glutathione (GGG) is a physiological ligand for P2RY8, which acts as a chemo-repellent that stops B cell migration in germinal centers^34^. Previous work has also shown that KO of P2RY8 in human T cells leads to enhanced transwell migration in the presence of GGG^35^. To test whether tumor cells may be secreting GGG, we used a previously described P2RY8 ligand bioassay that measures migration inhibition activity of culture supernatants on a mouse B lymphoma cell line (WEHI-231, naturally negative for P2RY8) transduced with a human P2RY8-GFP construct^34^. Here WT WEHI-231 cells are mixed with P2RY8^+^ WEHI-231 cells in a migration assay in the presence of culture supernatants and the chemoattractant chemokine CXCL12. Using this transwell bioassay, our data indicate that supernatants from all four tested human tumor cell lines can suppress migration in P2RY8+ WEHI-231 cells, but not in P2RY8 negative WT WEHI-231 cells and thus may contain GGG (**Fig. 3d, Extended Data Fig. 4b**). We further tested P2RY8 KO human T cells in the transwell assay, where P2RY8 KO and AAVS1 edited control T cells were mixed in a 50/50 ratio and then added to the transwell. We saw an increase in migration for P2RY8 KO human T cells compared to controls when tumor cell supernatant and CXCL12 were added to the bottom of the transwell, further supporting a role for P2RY8 in suppressing T cell chemotaxis toward cancer cells (**Fig. 3e and Extended Data Fig. 4c**). Additionally, we performed Liquid Chromatography with tandem mass spectrometry (LC-MS-MS) on multiple human cancer cell lines, and found that they all secrete GGG to varying degrees, suggesting that GGG produced by tumor cells could be regulating T cell entry via P2RY8 ligation (**Extended Data Fig. 4d**).

To directly assess the role of P2RY8 signaling in T cell migration into these tumors, we used an in vivo competition system. We transduced CD19 CAR T cells, NY-ESO-1 TCR T cells, or Claudin18.2 (CLDN18.2) CAR T cells with lentivirus encoding an sgRNA targeting either P2RY8 or control AAVS1, each linked to a unique fluorophore (GFP or BFP). These engineered T cells were mixed at a 50:50 ratio of P2RY8 KO (BFP) and AAVS1 control (GFP), and then injected into NSG mice bearing subcutaneous tumors with the cognate antigen (CD19 CAR T cells with CD19+ A375 melanoma cells, NY-ESO-1 TCR T cells with WT A375 melanoma cells that naturally express NY-ESO-1 and matched MHC-I, and Claudin 18.2 CAR T cells with OE19 gastroesophageal cancer cells that naturally express Claudin18.2). After 48 hours (an early time point chosen to measure effects of trafficking rather than proliferation), tumors and spleens were removed and T cells isolated to assess the P2RY8 KO to AAVS1 ratios. While the original 50:50 ratio was generally preserved in spleens, the P2RY8 KO T cells were significantly enriched compared to AAVS1 in the tumors, suggesting that P2RY8 may be inhibiting T cell infiltration into tumors in these models (**Fig. 3f**).

To assess the effect of P2RY8 KO on tumor control, we used CD19-specific CAR T cells with three solid tumor models all engineered to express CD19: a non-small cell lung cancer (NSCLC) cell line (A549), a pancreatic cancer cell line (AsPC-1), and a melanoma cell line (A375), as well as NY-ESO-1 TCR T cells with A375. These cell lines were subcutaneously injected into NSG mice, which were then treated with the relevant CAR T or TCR T cells edited for either P2RY8 or control AAVS1. On first testing in a fast-growing A375 tumor model, there was no appreciable advantage to the P2RY8 KO therapeutic T cells in tumor control (**Fig. 3g, Extended Data Fig. 4e-f**). However, testing in slower growing tumor models (A549 and AsPC-1) did show a trend towards slowed tumor growth. (**Fig. 3h-i**). We found evidence of higher accumulation of P2RY8 KO T cells in tumors relative to the AAVS1-targetd control T cells, but found no marked reduction in markers of exhaustion or dysfunction in the P2RY8 KO T cells (**Extended Data Fig. 4g**). These data suggest that while manipulation of cell trafficking pathways may enhance infiltration, which is an important step to improve tumor control, the T cells likely remain vulnerable to suppression and/or exhaustion within tumors, likely explaining the modest improvement in tumor control in our models. Based on this result, we hypothesized that sorting T cells by markers of their functional status at the endpoint of our in vivo screen could reveal gene targets that promote resistance to dysfunction within solid tumors.

### In vivo genome-wide screen identifies gene targets to prevent T cell dysfunction in solid tumors

To select for functional effector T cells in the TME, we isolated T cells from the tumors using the same approach as the tumor infiltration screens, and then sorted them based on interferon-gamma (IFN-γ) production. sgRNAs from the highest and lowest 20% of IFN-γ expressing TILs were sequenced to identify sgRNAs enriched in the high compared to the low IFN-γ expressing populations (**Fig. 4a and Extended Data Fig. 5a**). The sgRNAs recovered from each sorted condition were uniformly distributed, indicating good library representation **(Extended Data Fig. 5b)**. GSEA identified significant dropout of genes involved in IFN-γ production in the IFN-γ^hi^ versus IFN-γ^lo^ comparison, supporting the validity of the screen through the expected loss of known IFN-γ pathway regulators **(Extended Data Fig. 5c)**. Encouragingly, these screens identified many genes that are already well-described targets of interest to enhance human T cell therapies. For instance, our screen hits included TNFAIP3 (A20), RASA2, DGKZ, and CBLB, which have been proposed as gene targets to enhance human T cell therapies based on preclinical data^6,8,19,36–38^ (**Fig. 4b, Extended Data Table 2**). There were also many genes that scored highly in this screen that had not been previously linked to human T cell function or anti-tumor immunity. We compared our in vivo IFN-γ screen data with a previously published in vitro CRISPRi screen for IFN-γ production in human T cells^39^. Using a cutoff of FDR < 0.25, this comparison revealed partial overlap between hits, for example, DGKZ and CBLB, which regulate stimulation-induced IFN-γ production. However, many of our hits were found only in the in vivo setting, suggesting that these genes may play distinct roles in promoting anti-tumor effector function specifically within the TME (**Fig. 4c**).

**Fig. 4.**
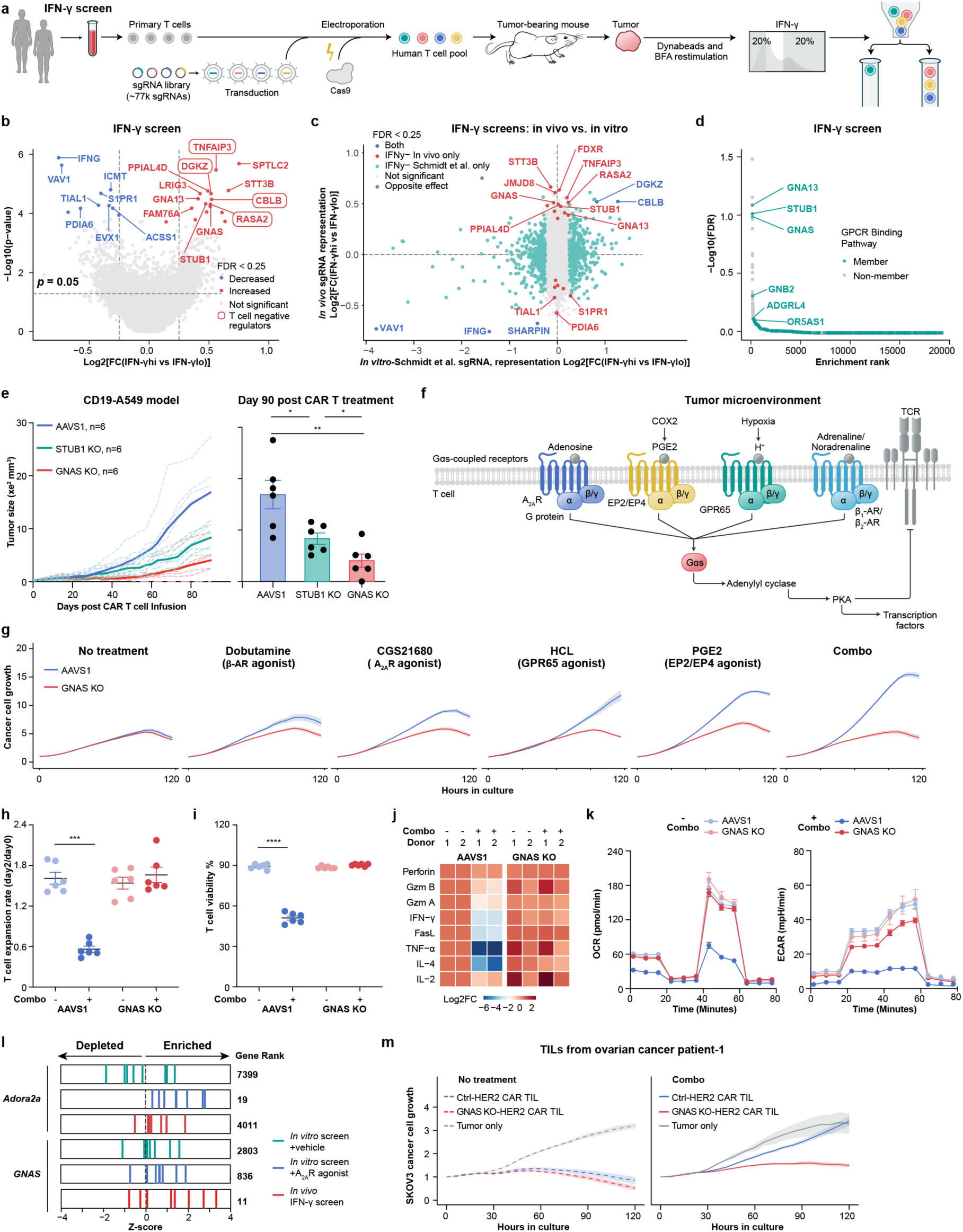
IFN-γ-based genome-wide in vivo CRISPR screen identifies key regulators of T cell function in solid tumors. **a.** Schematic of the in vivo genome-wide CRISPR IFN-γ screen in tumor infiltrating T cells. **b.** Volcano plots showing enriched (red) and depleted (blue) gene candidates associated with IFN-γ expression in TILs, as identified by MAGeCK analysis comparing IFN-γ high vs. low populations. The screen was performed using T cells from two human donors. Genes meeting the significance threshold (FDR < 0.25 and abs (Log2FC)≥ 0.25) are highlighted. Boxed genes represent known negative regulators of human T cell function. **c.** Comparison of primary human T cell IFN-γ screens: in vivo screen versus published in vitro screen from Schmidt et al^39^, colored by direction and specificity of effect between the two screens. Genes meeting the significance threshold (FDR < 0.25 and abs (Log2FC)≥ 0.25) are colored. **d.** Rank plot of the in vivo IFN-γ screen hits colored by gene association with GPCR binding pathways. The pathway annotation in this figure specifically includes genes were selected based on their association with the GPCR pathway from literature as well as genes in the msigdbr gene list. **e.** Tumor growth of CD19-A549 tumor-bearing NSG mice treated with CD19 CAR T cells 9 days after tumor injection (left) and tumor size at day 90 post CAR T cell infusion (right) (n = 6 mice for each group). **f.** Schematic of selected GPCR(s)-Gαs signaling pathways in T cells in the solid tumor microenvironment. Four of the known suppressive ligands found in the TME that bind GPCRs upstream of Gαs are indicated (schematic is representative, not comprehensive). **g.** Incucyte-based killing assay comparing GNAS KO and AAVS1 control CD19 CAR T cells against CD19 expressing A375 tumor cells, in the presence or absence of the indicated agonist or combination of all four agonists. Data are representative of one of three donors. **h-k.** AAVS1 and GNAS KO T cells were treated with a combination of the four agonists shown in (**g**) for 2 days in the presence of CD3/CD28 activation and in the absence of IL-2. Assessed outcomes include: T cell expansion rate on day 2 relative to day 0 (day 0 = day of bead stimulation) (**h**), cell viability on day 2 (**i**), log_2_ fold change (day 2/day 0) of cytokine secretion in culture supernatant (**j**), and metabolic activity on day 2 (**k**). n = 2 donors for h-j, and data are representative of one of three donors for k. **l.** sgRNA z-scores for selected genes of interest and gene rank from the in vivo IFN-γ screen and previously published in vitro screens treated with vehicle or Adora2a agonist, as reported by Shifrut et al, *Cell*, 2019. **m.** Incucyte-based killing assay comparing GNAS KO and AAVS1 control HER2 CAR TILs in the presence or absence of the four-agonist combination shown in (g). TILs were isolated from the ascites of an ovarian cancer patient and engineered to express HER2 CAR (other patient replicate shown in Extended Data Fig 5i). *P* values were determined by two-tailed unpaired Student’s t-test (**e, h, i**). \**P* < 0.05, \*\**P* < 0.01, \*\*\**P* < 0.001, and \*\*\*\**P* < 0.0001. Data are presented as mean ± s.e.m.

Gene set enrichment analysis showed enrichment for GPCR signaling mediators in the IFN-γ high T cells recovered from tumors (**Fig. 4d**). Top screen hits with roles in GPCR signaling pathways included GNAS, STUB1, and GNA13. GNA13 is part of the P2RY8 pathway (**Fig. 3b**), whereas GNAS and STUB1 are unique hits from the in vivo IFN-γ screen that were not identified in the previously published in vitro IFN-γ screen. These genes did not drop out in the tumor abundance screen (**Extended Data Fig. 5d**), in contrast to other unique hits from the in vivo IFN-γ screen, such as STT3B, FDXR, JMJD8, and PPIAL4D (**Fig. 4c**). We therefore chose these two uniquely in vivo enriched gene targets to test if they enhanced clearance in our CD19-expressing A549 slow-growing NSCLC tumor model. GNAS and STUB1 KO CD19 CAR T cells were injected alongside AAVS1 control-edited CAR T cells into A549 tumor-bearing NSG mice. Although knockout of both genes improved tumor control, the striking potency of GNAS KO CAR T cells led us to prioritize this target for in-depth functional analysis (**Fig. 4e, Extended Data Fig. 5e, f**).

GNAS (ranked 11 in our in vivo IFN-γ production screen) encodes Gαs, the stimulatory alpha subunit of a heterotrimeric G protein complex that couples with GPCRs. Upon GPCR activation, Gαs dissociates from the β and γ subunits and activates adenylyl cyclase, leading to the production of cAMP, activation of Protein Kinase A (PKA), and PKA-mediated phosphorylation of multiple targets involved in metabolic regulation, transcription, and signal transduction^40^. Gαs can function as a central mediator of signaling from multiple T cell GPCRs that sense a variety of suppressive ligands^41^ (**Fig. 4f**). For instance, Gαs can be activated by GPCRs such as ADORA2A (adenosine receptor), EP2/EP4 (receptors for prostaglandin), GPR65 (acidity (H^+^) receptor), and β1-and β2-adrenergic receptors (receptors for adrenaline and noradrenaline), among others^21,41–47^. These GPCRs serve as receptors for suppressive ligands found in the TME that can induce T cell dysfunction. Our screen data suggest that genetic ablation of GNAS in human T cells can program them to ignore suppressive cues in the TME and thereby enhance their intra-tumoral effector function. To test whether disruption of GNAS enhances T cell resistance to suppressive cues characteristic of the tumor microenvironment, we treated co-cultures with individual or combined suppressive factors, including prostaglandin E₂ (PGE₂), an adenosine A₂A receptor agonist (CGS21680), acidic pH (HCl), and dobutamine, and assessed tumor cell killing in vitro. Interestingly, in the absence of suppressive ligands, GNAS KO CD19 CAR T cells showed no significant advantage over AAVS1 control T cells in tumor cell killing. However, upon addition of individual suppressive cues, the GNAS KO T cells exhibited varying degrees of enhanced cytotoxicity (**Fig. 4g**). Notably, when all suppressive ligands were combined, the advantage conferred by GNAS disruption became particularly pronounced. We also measured T cell expansion and viability in the presence or absence of these suppressive treatments and confirmed that while the AAVS1 control T cells were potently suppressed, the GNAS KO T cells appeared unaffected (**Fig. 4h-i**). Similarly, compared to control T cells, GNAS KO T cells maintained robust expression of activation markers and sustained production of cytokines and effector molecules, remaining largely resistant to the combined suppressive effects of the ligands in vitro (**Fig. 4j, Extended Data Fig. 5g**). Given that metabolic function is central to persistent T cell function in the TME^4^, we tested the effects of these suppressive ligands on T cell metabolism using an extracellular flux assay for real-time measurement of cellular metabolism (Seahorse assay). We found that two parameters, the oxygen consumption rate (OCR) used to determine aerobic mitochondrial respiration, and the extracellular acidification rate (ECAR), used to measure anaerobic glycolysis, were highly resistant to the suppressive effects of the combined suppressive ligands when GNAS was knocked out (**Fig. 4k, Extended Data Fig. 5h**).

These data suggest that Gαs is a central integrator of multiple suppressive cues that can disable T cells within tumors. We reviewed data from a prior in vitro genome-wide CRISPR KO screen, where we challenged T cells with either vehicle or with the adenosine (A2A) receptor agonist, CGS21680 and sorted the cells based on cell division dye levels to isolate the more proliferative cells. Interestingly, sgRNAs against GNAS were modestly enriched in the CGS21680 treated compared to vehicle arms, corroborating its role in adenosine receptor signaling (**Fig. 4l**). However, the rank score of GNAS in this in vitro screen (928) was far less robust compared to its rank score of 11 in our in vivo screen. The relatively modest advantage observed for GNAS knockout in the in vitro screen parallels its limited benefit in the presence of individual suppressive ligands (**Fig. 4g**), an effect that is markedly amplified when these ligands are combined. This contrast underscores a key advantage of performing screens in vivo, where gene perturbations such as GNAS disruption are selectively enriched under conditions that impose multiple, overlapping suppressive pressures, thereby revealing targets with broader and more physiologically relevant impact.

Our data demonstrate that GNAS KO prevents T cell dysfunction in healthy donor T cells. However, since therapeutic application would involve gene editing of patient-derived T cells, either peripheral blood or tumor-infiltrating lymphocytes (TILs), which have already been exposed to systemic and intratumoral cues that induce dysfunction, we sought to evaluate whether GNAS KO could also reverse or prevent dysfunction in patient TIL. To this end, we transduced TIL isolated from patients with ovarian cancer with a HER2-specific CAR and edited them to target either AAVS1 (control) or GNAS. These CAR TILs were then co-cultured with HER2⁺ tumor cells, with or without the cocktail of combined suppressive ligands. Consistent with our findings in healthy donor T cells, GNAS KO conferred remarkable resistance to cytolytic dysfunction in patient-derived TILs under suppressive conditions (**Fig. 4m, Extended Data Fig. 5i**), indicating that GNAS disruption can preserve the function of patient TILs even in hostile, tumor-like environments.

The disparity of effect size when comparing GNAS KO cells in the presence of single vs. combined suppressive ligands suggests that Gαs is functioning as a central integrator of various suppressive cues that antagonize anti-tumor T cell function. We assessed sgRNA-level enrichment for GPCRs that recognize the key suppressive ligands tested, across both our IFN-γ and abundance screens. We noted that while many of these GPCRs had 2-3 sgRNAs with high positive LFC levels, GNAS (IFN-γ screen) and PTGER4 (abundance screen) clearly stood out, ranking among the top hits (**Extended Data Fig. 6a**). To examine this further, we assessed knockout of individual GPCRs upstream of GNAS in an attempt to phenocopy such partial functional enhancements. We found that indeed, knockout of individual GPCRs can boost tumor control, but knockout of GNAS has a much more potent effect as it simultaneously integrates the benefits of losing multiple GPCRs together (**Extended Data Fig. 6b**). These data underscore the value of conducting functional screens for enhanced anti-tumor T cell activity in physiologically relevant in vivo settings. They also highlight the therapeutic potential of identifying convergent regulators that integrate multiple suppressive signals within the tumor microenvironment, enabling more robust T cell function than targeting individual inhibitory pathways alone.

### GNAS KO in therapeutic human T cells potently enhances efficacy in multiple preclinical solid tumor models

We next tested the effects of disrupting GNAS in therapeutic T cells in a large array of solid tumor models. We subcutaneously injected NSG mice with CD19-expressing A375 tumor cells, and 9 days later injected CD19 CAR T cells from two different donors, edited with sgRNAs for either AAVS1 or GNAS. While only 1 out of 6 mice receiving AAVS1 CAR T cells was able to control tumor growth at this dose, all 6 of the mice receiving GNAS KO CAR T cells experienced complete and durable tumor clearance, corresponding to a dramatic survival benefit (**Fig. 5a**). The results were similarly striking using a second T cell donor (**Extended Data Fig. 7a-b).** In order to test for effects of GNAS KO on T cell memory and persistence, we rechallenged mice that received GNAS CAR T cells with a second tumor injection 59 days after the first (**Fig. 5b**). Specifically, these mice were injected with CD19-expressing A375 tumor cells in one flank, and CD19-negative A375 tumor cells in the opposite flank to evaluate whether GNAS disruption can enhance functional persistence and rejection of the antigen-positive tumors. GNAS KO CAR T cells demonstrated robust anti-tumor memory in this rechallenge model, as indicated by sustained rejection of antigen-positive tumors, whereas antigen-negative tumors progressed unimpeded (**Fig 5b**). Flow-based analyses of these tumors at the end of the experiment showed a trend towards higher CAR T cell infiltration in the antigen-positive compared to the antigen-negative tumors (**Extended Data Fig. 7c**). In order to recover GNAS KO vs AAVS1 control CAR T cells from the initial tumors for further characterization, we repeated this experiment with a lower dose of CD19 CAR T cells such that we would not see complete tumor clearance (**Fig. 5c, Extended Data Fig. 7d**). There was a dramatic increase (∼20 fold) in the average number of GNAS KO CAR T cells per gram of tumor tissue measured by flow cytometry (**Fig. 5d, Extended Data Fig. 7e**). Additionally, the GNAS KO CAR T cells were markedly less exhausted based on multiple canonical exhaustion markers, with CD39 showing a nearly 10-fold lower MFI (**Fig. 5e**). Thus, we observed a striking difference in both the frequency and functional state of GNAS KO CAR T cells within tumors, despite minimal differences in their intrinsic fitness observed in vitro (**Fig. 4g-j, Extended Data Fig. 5g-h**).

**Fig. 5.**
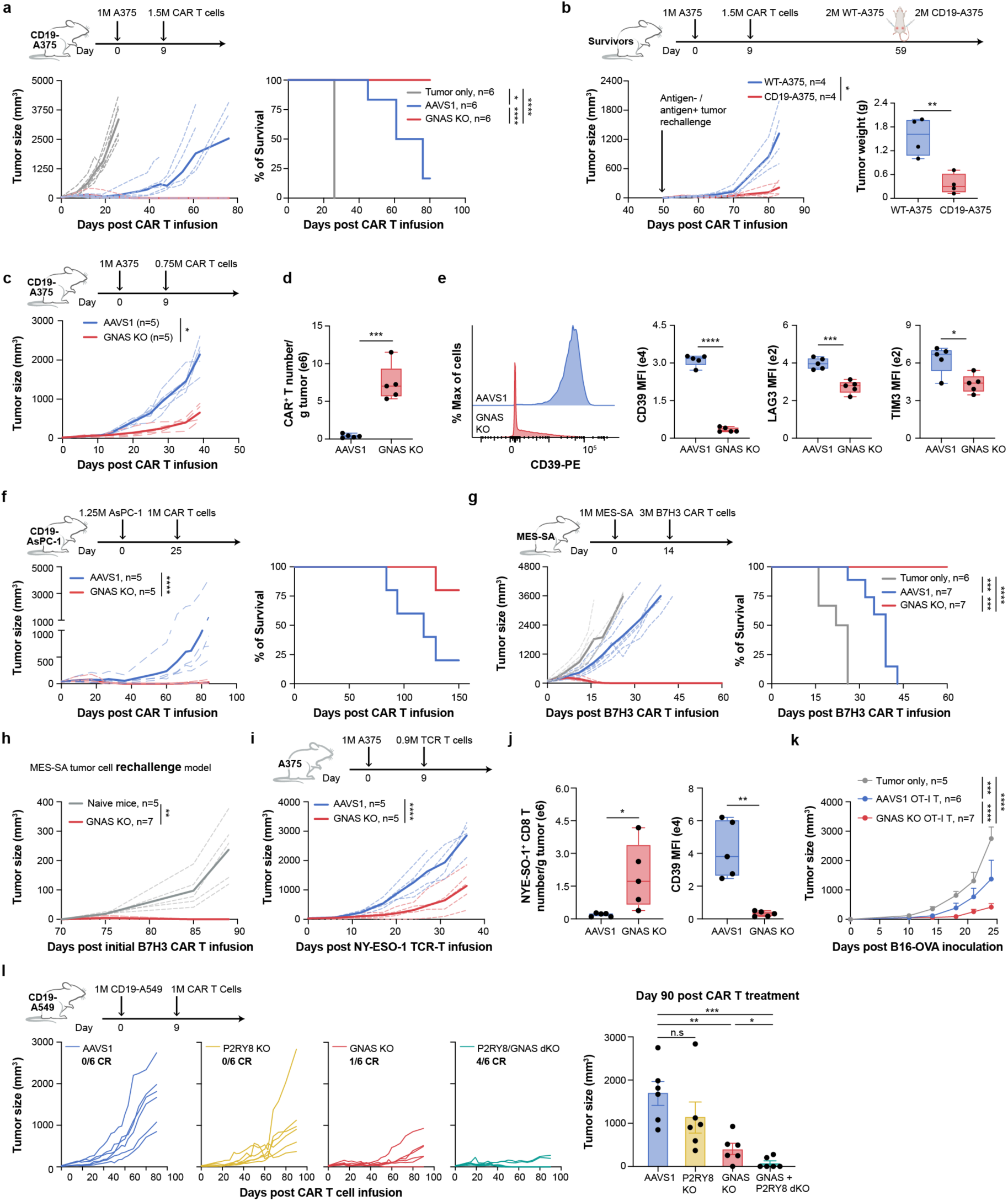
GNAS ablation enhances CAR T cell fitness in multiple preclinical solid tumor models. **a.** CD19-A375 melanoma cells were subcutaneously engrafted into NSG mice, followed by intravenous infusion of CD19 CAR T cells. The experimental timeline (top), tumor growth curves (bottom left), and mouse survival (bottom right) are shown (n = 6 mice per group). Data are from one of two donors, data from the other donor shown in Extended Data Fig.7b. **b.** Surviving mice treated with the GNAS KO CAR T cells from (a) that had complete responses were rechallenged with new tumor cells. Antigen negative wildtype A375 and antigen positive CD19-A375 tumor cells were subcutaneously engrafted into each flank on day 59 post-original tumor challenge. Tumor growth (bottom left) and tumor weights (bottom right) are shown (n = 4 mice). **c-e.** CD19-A375 melanoma cells were subcutaneously engrafted into NSG mice, followed by intravenous infusion of a low dose of CD19 CAR T cells to enable in vivo T cell isolation and phenotypic analysis. The experimental timeline (**c**, top), tumor growth curves (**c**, bottom), absolute number of CAR T cells per gram of tumor (**d**), and expression levels of CD39, LAG-3, and TIM-3 on tumor-infiltrating CAR T cells (**e**) are shown at day 39 post T cell infusion (n = 5 mice for each group). **f.** CD19-AsPC-1 pancreatic cancer cells were subcutaneously engrafted into NSG mice, followed by intravenous infusion of CD19 CAR T cells. The experimental timeline (top), tumor growth curves (bottom left), and mouse survival (bottom right) are shown (n = 5 mice for each group). **g.** MES-SA uterine sarcoma cells were subcutaneously engrafted into NSG mice, followed by intravenous infusion of B7H3 CAR T cells. The experimental timeline (top), tumor growth curves (bottom left), and mouse survival (bottom right) are shown (n = 6 mice for the tumor only control group, and n = 7 mice for both the AAVS1 control and GNAS KO CAR T cell groups). **h.** MES-SA tumor growth upon tumor rechallenges. Naïve NSG mice were newly challenged and surviving mice from (g) that experienced complete responses were rechallenged with MES-SA tumor cells on day 70 post initial B7H3 CAR T cell infusion. n = 5 for naïve mice and n = 7 for GNAS KO surviving mice from (g). **i-j.** A375 melanoma cells were subcutaneously engrafted into NSG mice, followed by intravenous infusion of NY-ESO-1 TCR T cells. The experimental timeline (**i**, top), tumor growth curves (**i**, bottom), absolute number of NY-ESO-1 specific CD8^+^ TCR cells per gram of tumor (**j**, left), and expression levels of CD39 tumor-infiltrating NY-ESO-1 specific CD8^+^ TCR cells (**j**, right) are shown at day 37 post T cell infusion (n = 5 mice for each group). This experiment includes data already shown in Extended Data Fig. 4e-f. All experimental conditions for these figure panels were performed together as a concurrent experiment and some plots are shown elsewhere for narrative clarity. **k.** Tumor growth curves following adoptive transfer of AAVS1 control or GNAS KO OT-I CD8⁺ T cells (7 × 10^6^) into C57BL/6 mice bearing subcutaneous B16-OVA tumors. OT-I T cells were administered on day 10 post-tumor engraftment (n = 5 mice for the no T cell control group, n = 6 mice for the AAVS1 OT-I T cell group, and n = 7 mice for the GNAS KO OT-I group). **l.** Tumor growth curves for mice treated with CD19 CAR T cells with individual or combined P2RY8 and GNAS gene KOs. CD19-expressing A549 non-small cell lung cancer cells were subcutaneously engrafted into NSG mice, followed by intravenous infusion of CD19 CAR T cells. The experimental timeline (top), tumor growth curves (bottom left), and tumor size at day 90 post CAR T cell treatment (bottom right) are shown (n = 6 mice per group). This experiment includes data already shown in Fig. 3h, Fig. 4e and Extended Data Fig. 6b. All experimental conditions for these figure panels were performed together as a concurrent experiment and some plots are shown elsewhere for narrative clarity. *P* values were determined by two-way ANOVA in (**a, b, c, f, g, h, i, k**) for tumor growth, Log-rank (Mantel-Cox) test in (**a, f, g**) for mice survival analysis, and two-tailed unpaired Student’s t-test in (**d, e, j, l**). \**P* < 0.05, \*\**P* < 0.01, \*\*\**P* < 0.001, and \*\*\*\**P* < 0.0001. n.s, not significant. Data are presented as mean ± s.e.m.

To address broader therapeutic potential, we tested the effects of GNAS KO in additional preclinical solid tumor models. We found that GNAS KO CD19 CAR T cells showed a robust advantage in controlling the growth of a pancreatic cancer model, AsPC-1 engineered to express CD19 (**Fig. 5f**). Next, we tested B7H3-specific CAR T cells^48^, which are currently in clinical trials for multiple solid tumor indications^49^, in NSG mice subcutaneously engrafted with a uterine sarcoma cell line (MES-SA, naturally expresses B7H3). Mice treated with AAVS1 control CAR T cells experienced rapid tumor progression, whereas GNAS KO B7H3-specific CAR T cells not only eradicated the initial MES-SA tumor but also conferred durable protection, rejecting a tumor rechallenge administered on day 70 after CAR T cell infusion (**Fig. 5g, h**). This enhanced tumor control was maintained even at lower CAR T cell doses (**Extended Data Fig. 7f-g**). We further tested effects of GNAS KO in TCR transgenic tumor models. NY-ESO-1 specific human 1G4 TCR T cells were injected into mice that were subcutaneously injected with A375 tumor cells, which naturally express and present NY-ESO-1 peptide on cognate MHC-I. In this model system, GNAS disruption conferred a clear tumor control advantage, accompanied by enhanced TCR T cell accumulation and reduced exhaustion in the tumor (**Fig. 5i-j**). A similar benefit was observed in murine GNAS KO OT-I T cells within a fully immunocompetent in vivo setting (**Fig. 5k**).

Given the clear therapeutic promise, we proceeded to test basic safety metrics, specifically whether GNAS KO CAR T cells remain cytokine and antigen dependent. As previously noted, there was no appreciable difference between the AAVS1 and GNAS KO T cells in terms of stimulation-dependent proliferation or tumor cell killing in vitro (**Fig. 4g-h**). GNAS KO CD19 CAR T cells demonstrated contraction and loss of viability as expected upon cytokine withdrawal, similar to AAVS1 control CAR T cells (**Extended Data Fig. 8a**). Additionally, we found no evidence of off-target killing by GNAS KO CD19 CAR T cells co-cultured with tumor cells negative for target antigen (**Extended Data Fig. 8b**). We monitored mice from our AsPC-1 and MES-SA models, which we engineered by targeted integration of CD19 CAR or B7H3 CAR at the TRAC locus^50^, and found no evidence of weight loss or signs of poor health in the mice receiving GNAS KO CAR T cells (**Extended Data Fig. 8c-f**). These results support a favorable preclinical safety profile for GNAS KO CAR T cells.

Finally, as our two in vivo screening readouts presumably highlighted gene targets that change T cell behaviors along unique and potentially complementary axes, we hypothesized that combining two gene edits in one cell product might lead to useful synergistic properties for tumor control. Our sgRNA abundance screen enabled the discovery that targeting the P2RY8-Gα13-ARHGEF1 signaling axis enhances tumor infiltration but that these cells remain susceptible to various forms of suppression inside the tumor. Our IFN-γ screen highlighted GNAS as a potent gene target to preserve T cell effector function by disabling suppressive signals within the tumor. We reasoned that we could combine disruption of P2RY8 and GNAS to simultaneously enhance trafficking and confer resistance to suppression, which could be synergistic for tumor control. We subcutaneously engrafted NSG mice with a CD19-expressing A549 NSCLC cell line, followed by intravenous infusion of CD19 CAR T cells edited at the AAVS1 control locus, P2RY8 KO, GNAS KO, or GNAS/P2RY8 dual KO. While the GNAS KO effect is quite strong, we found that combining with P2RY8 KO led to an even greater improvement in overall tumor control, with more mice receiving the dual edited CAR T cells achieving complete tumor clearance (AAVS1: 0/6 complete responses (CRs), P2RY8 KO: 0/6 CR, GNAS KO: 1/6 CR, GNAS + P2RY8 KO: 4/6 CR) (**Fig. 5l**). Together, these results suggest that our in vivo screens uniquely identified genetic modifications that enhance T cell trafficking into tumors and T cell resistance to immunosuppression in the TME, which can be combined to achieve more effective CAR T cell products with significant potential to treat solid tumors.

## Discussion

In this study, we developed a novel screening approach that enables, for the first time, genome-scale CRISPR screens in human T cells in solid tumor-bearing mice. This new strategy offers a solution to the relative limitations of in vitro screens, that do not model many of the complexities of the intratumoral challenges experienced by T cells in a living organism. By moving the discovery work in vivo, we can accelerate prioritization of gene engineering enhancements for T cell therapies to treat patients with solid tumors. While our screening system is mechanically most similar to a TIL-based model, we show that the targets nominated by our screens are relevant in engineered TCR and CAR T cell therapies in a large range of preclinical solid tumor models. Our abundance and effector state screens highlighted the GPCR signaling pathways, P2RY8-Gα13-ARHGEF1 and GPCR(s)-Gαs-PKA, respectively, as key pathways governing T cell entry and efficacy in solid tumors. Given the solid tumor biology underlying these effects, in vitro screening systems would be unlikely to uncover such pathways affecting T cell trafficking or resistance to combined suppressive signals.

Our discovery that P2RY8 receptor signaling is a potential regulator of immune exclusion co-opted by tumors showcases the importance of a screening system directly within primary human T cells in tumor-bearing animals. While many vertebrates have orthologues of P2RY8, rodents lack an orthologue of this gene, and thus it would not have been uncovered in a fully murine in vivo screen. This highlights a strength of our screening system, which can uncover functional roles for human genes in human T cells in tumor-bearing mice. Tumors may be co-opting the P2RY8 signaling pathway, which is important for lymphocyte trafficking, to block T cell migration and reduce infiltration. There have been a few examples of GPCR ligands acting directly on T cells to establish immune exclusion. For instance, CXCL12 can be secreted by cancer-associated fibroblasts to act as a retention signal to trap CXCR4-expressing T cells in the stromal regions away from the cancer cells^31^. Additionally, endothelial cells in the tumor can express Semaphorin 3A, which binds to NRP1 and plexin receptors on T cells, inducing T cell paralysis and impairing migration into the tumor core^51^. The Gα13-coupled lysophosphatidic acid receptor, LPAR6 (also known as P2RY5), has also been suggested to contribute to tumor immune exclusion based on inhibitor studies, though this finding has not yet been confirmed with genetic knockout experiments^52^. Here, we have uncovered another putative mechanism of immune exclusion, whereby tumor cells secrete ligands, such as GGG, to reduce T cell entry. While full characterization of the role of this pathway in human T cell migration and growth in tumors will require further investigation, this finding demonstrates how unbiased screening in human T cells *in vivo* can uncover novel biology that previously would have been inaccessible.

We identified multiple GPCRs and their signaling components that serve to restrain T cell production of IFN-γ in the in vivo tumor environment, of which we focused on GNAS (Gαs). Ablation of GNAS disables a central signaling node that integrates suppressive cues from several upstream GPCRs. Previously, Wu et al. used a transgenic mouse model expressing a CD8-restricted Gαs–DREADD to activate CD8-restricted Gαs signaling and demonstrated that activation of the Gαs–PKA signaling axis promotes CD8^+^ T cell dysfunction and resistance to immune checkpoint inhibitors^41^. In addition, prior studies have focused on disabling a key downstream mediator of Gαs signaling, PKA, in T cells by developing a synthetic peptide that disrupts ezrin–PKA association, called the “regulatory subunit I anchoring disruptor” (RIAD)^53,54^. They showed that expression of RIAD displaces PKA from lipid rafts and diminishes PKA’s phosphorylation of Y505 on Lck, leading to upregulated TCR signaling, and that expression of RIAD in CAR T cells can enhance tumor killing in mouse models^54^. Thus, while prior studies in other contexts have implicated Gαs signaling in T cell dysfunction, this signaling node has not been previously targeted in human T cells to enhance cancer immunotherapies. Our advanced in vivo screening platform nominated GNAS as a potent gene editing target to overcome human T cell therapy failure in solid tumor environments.

Taken together, in addition to uncovering novel biology, including a role for P2RY8 in restraining human T cell trafficking into tumors, our screening system was able to accurately and rapidly prioritize gene targets for immediate testing in cell therapy products for solid tumors. We find that a single KO edit of GNAS leads to dramatic improvements in solid tumor control across multiple preclinical models. The fact that we see only meager to no effects of GNAS deletion on T cell function in vitro reinforces that our screen was able to identify a gene target that selectively enhances T cell function in the solid tumor environment. Indeed, our experiments with combined suppressive ligands reveal that the full benefit of GNAS knockout only becomes apparent when multiple suppressive cues are present. Excitingly, our complementary screens allowed us to identify a combinatorial approach to program therapeutic cells, combining GNAS KO with P2RY8 KO to enhance both trafficking and resistance to suppression in T cell products. This platform provides an adaptable screening framework to identify synergistic gene perturbations that can be multiplexed to enhance T cell efficacy. Given the complexity of the tumor microenvironment, a multiplex gene editing strategy will likely be necessary to endow T cells with the capacity to overcome these multifaceted challenges. Our initial screens using this model underscore its unique utility in uncovering diverse, therapeutically relevant approaches to optimize T cell therapies for solid tumors.

The system described here enables genome-scale discovery in human T cells within tumor-bearing animals, capturing key complexities of the solid tumor microenvironment that are absent in traditional in vitro models. Notably, in comparison to in vitro systems, this platform should better recapitulate biophysical and biochemical barriers to T cell infiltration, intensified competition for nutrients and oxygen, accumulation of suppressive metabolites, and the integration of multiple inhibitory cues. The concurrent exposure of T cells to these diverse pressures likely constitutes a major advantage of in vivo screening, allowing the identification of genetic perturbations that confer context-specific fitness. However, this model is not immunocompetent and lacks several immunosuppressive components of the tumor microenvironment, including regulatory T cells, myeloid-derived suppressor cells, tumor-associated macrophages, and the cytokine milieu produced by an intact immune system. Future iterations of this system could incorporate recent advances in humanized mouse models, such as NSG-SGM3 and related strains, that support multilineage human hematopoiesis through expression of human cytokines enabling engraftment with hematopoietic stem cells or PBMCs^55–57^. Moreover, this platform could be adapted to fully immunocompetent murine systems by leveraging a human/mouse CD3ε chimera strategy. Endogenous mouse T cells do not bind OKT3 and therefore would not reject tumors engineered to express the OKT3 scFv, facilitating stable tumor engraftment. Murine T cells from a previously reported mouse model expressing a human/mouse CD3ε chimera^58^, could then be harvested and used for pooled in vivo screening, as they express the human CD3ε epitope recognized by the OKT3 scFv and would thus engage the tumors upon transfer. This approach would enable large-scale in vivo screening in mouse T cells within fully immunocompetent, tumor-bearing hosts. We envision multiple such strategies to further increase the physiological relevance of our screening platform and more accurately model the complexity of the immunocompetent tumor microenvironment.

The discovery potential of our screening approach goes well-beyond the screens demonstrated in this study. This system should be highly flexible and adaptable to a variety of screening formats, including libraries of pooled sgRNAs, transgenes, or other constructs that can be introduced into T cells. For example, it could be used to identify gain-of-function genes or gene variants that enhance T cell function in vivo, through either CRISPR activation^37^ or expression of ORF libraries. The scalability of this platform will be of high utility to perform mutagenesis screens using base editing libraries for many genes simultaneously, enabling exploration of key domains and residues within genes of interest to tune gene functions in vivo. This platform would also be capable of screening large libraries for multiplexed discovery, using tools such as Cas12a or Cas13d^59,60^ to identify synergistic gene combinations. There is also great interest emerging for screening synthetic genes, such as switch receptors or genes with combinations of shuffled domains^61^, and this platform could be used to rapidly assess their effects on T cell performance in vivo. In addition, this model could be adapted to any tumor cell line, allowing in vivo testing against multiple tumor types. While this study focused on total abundance and IFN-γ production as screening readouts, the endpoint is highly customizable: T cells can be sorted based on functional surface markers of interest or linked to single-cell transcriptional states via Perturb-seq. Finally, in this work we used healthy human donor T cells; however, it would be feasible to screen TIL from cancer patients in this system to identify genetic enhancements that can reinvigorate TIL that have already been epigenetically rewired into exhausted states. Together, this platform represents a promising and versatile strategy for accelerating the discovery of next-generation genetic engineering approaches to improve T cell efficacy against solid tumors.

## Methods

### Lentiviral plasmid cloning

The anti-CD3 scFv construct was generated using a lentiviral vector backbone (Addgene #129443). This backbone was assembled with a gBlock containing a CD8 signal peptide, an anti-CD3 variable heavy chain (VH), a flexible linker, an anti-CD3 variable light chain (VL), a CD8 hinge region, a CD8 transmembrane domain, and a FLAG tag, all driven by the EF-1α core promoter on the backbone. The VH and VL sequences were derived from the commercial CD3 monoclonal antibody OKT3. For sgRNA expression, gBlocks ordered from IDT were cloned into either the LRG2.1-TagBFP2 (Addgene #124773) or LRG2.1-EGFP (Addgene #108098) lentiviral backbones. The backbones were digested with BsmBI-v2, and gBlocks were inserted using Golden Gate Assembly following the manufacturer’s instructions.

### Human T cell isolation and culture

Human peripheral blood leukopaks from anonymous healthy donors were obtained from Stemcell Technologies (catalog no. 200-0092) under Institutional Review Board–approved protocols. CD3⁺ primary human T cells were isolated using the EasySep™ Human T Cell Isolation Kit (catalog no. 17951) following the manufacturer’s instructions. Immediately after isolation, T cells were either cryopreserved or used directly in downstream experiments. Freshly isolated T cells were seeded at 1 × 10⁶ cells/mL in appropriate culture vessels and activated using CST Dynabeads (Thermo Fisher Scientific, 40203D) at a 1:1 bead-to-cell ratio. Unless otherwise specified, cells were cultured in cX-VIVO medium: X-VIVO™ 15 (Lonza Bioscience, 04-418Q) supplemented with 5% fetal bovine serum (Fisher Scientific), 50 µM 2-mercaptoethanol (Thermo Fisher Scientific, 21985023), and 10 mM N-acetyl-L-cysteine (Neta Scientific, SIAL-A7250-50G) in the present of 50U human IL-2 from R&D systems (202-GMP-01M) or 30U human IL-2 from NIH (BULK Ro 23-6019). For cryopreservation, T cells were frozen in RPMI-1640 medium containing 20% fetal bovine serum and 10% dimethyl sulfoxide at a concentration of 20–50 million cells/mL. Where indicated, GPCR agonists were added at the following concentrations: 1 μg/mL prostaglandin E₂ (Thomas Scientific, C956T61), 10 μM dobutamine hydrochloride (Tocris, 0515), 20 μM CGS-21680 hydrochloride (Tocris, 1063), and 10 mM HCl (adjusted to pH 6.6 in the final medium).

### Viral production and human T cell transduction

For lentiviral production, 27-28 × 10⁶ Lenti-X HEK 293T cells were seeded per poly-L-lysine– coated T225 flask in 45 ml of complete Opti-MEM (cOpti-MEM; Opti-MEM supplemented with 5% FBS and 1% penicillin-streptomycin, 1x NEAA, 1mM sodium pyruvate) and cultured overnight. Cells were transfected with 36 μg of the plasmid of interest, 36 μg psPAX2 (Addgene #12260), and 15 μg pMD2.G (Addgene #12259) using 144 μl P3000 and 168 μl Lipofectamine 3000 (Fisher Scientific, L3000075). Six hours post-transfection, the medium was replaced with fresh cOpti-MEM supplemented with 1× Viral Boost (Alstem, VB100). Virus-containing supernatants were collected at 18 hours and 42 hours post-media change, spun at 500 × g for 10 minutes at 4 °C to remove debris, and concentrated using Lenti-X Concentrator (Takara Bio, 631232) following the manufacturer’s instructions. The concentrated virus was stored at-80 °C until use. For T cell transduction, 24 hours after Dynabeads activation, the concentrated lentivirus was added directly to T cells at a 1:50 (v/v) ratio with X-Vivo-15 medium and gently mixed by tilting.

For AAV production, HEK293T cells were co-transfected with the AAV cargo plasmid containing the HDR template for the TRAC CAR construct and packaging plasmids to generate AAV6 particles. Viral particles were purified by iodixanol gradient ultracentrifugation. Viral titers were quantified by quantitative PCR (qPCR) following DNase I (NEB) treatment and proteinase K (Qiagen) digestion of the AAV preparations.

### Cell lines

The cell lines used in this study included A375 melanoma cells (ATCC, CRL-1619), A549 lung epithelial carcinoma cells (ATCC, CCL-185), AsPC-1 pancreatic adenocarcinoma cells (ATCC, CRL-1682), OE19 human Caucasian esophageal carcinoma cells (Sigma, 96071721-1VL), MES-SA uterine sarcoma cells (ATCC, CRL-1976), SKOV3 ovarian cancer cells (ATCC, HTB-77), B16-OVA MO4 Mouse Melanoma Cell Line (Sigma, SCC420) and Lenti-X HEK 293T cells (Takara Bio, catalog no. 632180). Anti-CD3 scFv or CD19-overexpressing A375 cells were generated in-house. MES-SA and CD19-overexpressing AsPC-1 cells were kindly provided by the Greg Allen lab at the University of California, San Francisco (UCSF). CD19-overexpressing A549 cells were gifts from the Kole Roybal lab at UCSF, and OE19 cells were provided by Alan Ashworth at UCSF. P2RY8 wild-type and P2RY8-expressing WEHI-231 cells were gifts from the Jason G. Cyster lab at UCSF. A375, AsPC-1, OE19 and SKOV3 cells were cultured in RPMI medium supplemented with 10% FBS, 50 IU penicillin/streptomycin, and 2 mM GlutaMAX. A549 and Lenti-X HEK 293T cells were cultured in DMEM with 10% FBS and penicillin/streptomycin. MES-SA cells were cultured in McCoy’s 5A Medium (ATCC, 30-2007) supplemented with 10% FBS and 50 IU penicillin/streptomycin. P2RY8-expressing WEHI-231 cells were cultured in RPMI containing 10% FBS, 10 mM HEPES, 2 mM glutamine, 55 μM 2-mercaptoethanol, and 50 IU penicillin/streptomycin. All cell lines were routinely tested for mycoplasma contamination by using Lonza Walkersville MycoAlert Mycoplasma Detection Kit (Fisher scientific, NC 9719283).

### In vivo screening and analysis

T cells isolated from two healthy human donors were stimulated as described above. After 24 hours, they were transduced with a genome-wide sgRNA library (Brunello library; Addgene) at a predefined titer to achieve approximately 50% transduction efficiency. Twenty-four hours later, the cells were washed with PBS and electroporated with Cas9 RNPs complexed with a non-targeting sgRNA (Horizon Discovery Biosciences Ltd., U-007503-01-20). Cells were then cultured and expanded in cX-VIVO medium supplemented with 50 U/mL IL-2 (R&D Systems, 202-GMP-01M) as previously described. At 72 hours post-transduction, 2 µg/mL puromycin was added to select for sgRNA⁺ T cells, and selection continued for six days. On day 11 post-isolation, 4 × 10⁶ transduced T cells were intravenously injected into NSG mice (n = 22 per donor), which had been engrafted with A375^low^ tumor cells 17 days prior. An aliquot of 4 × 10⁷ transduced T cells was reserved as the input sample (∼500× sgRNA coverage). Ten days post-infusion, mice were sacrificed, and tumors and spleens were harvested. Tumors were enzymatically digested, and T cells were isolated using Ficoll density gradient centrifugation; spleens were dissociated mechanically using syringe plungers. For each donor, tumor-infiltrating and splenic T cells were pooled separately. Final yields ranged from 190–230 million TILs and 100–150 million splenic T cells per donor. Genomic DNA (gDNA) was extracted using a phenol:chloroform:isoamyl alcohol protocol. For every 5 × 10⁶ T cells, cell pellets were thawed and incubated overnight at 65 °C in 400 µL Chip lysis buffer containing 16 µL of 5 M NaCl. The following day, 32 µL RNase A (10 mg/mL, QIAGEN, 19101) was added and incubated at 37 °C for 2 hours, followed by the addition of 16 µL proteinase K (20 mg/mL, Promega, V3021) and incubation at 55 °C for 2 more hours. gDNA was then purified by extraction with phenol:chloroform:isoamyl alcohol (Sigma-Aldrich, P3803-400ML), precipitated with sodium acetate, and washed with 70% ethanol. DNA was eluted in water and quantified using the 1X dsDNA High Sensitivity Assay Kit (Thermo Fisher) on a Qubit Fluorometer (Invitrogen). For abundance screens, gDNA was extracted from splenic T cells and from half of the tumor-infiltrating T cells. sgRNA sequences were PCR-amplified using Ex Taq DNA Polymerase (Takara Bio) with P5/P7 primers (IDT) following a standard protocol (https://portals.broadinstitute.org/gpp/public/dir/download?dirpath=protocols/production&filename=sgRNA_PCR_for_Illumina_sept2015.pdf), purified using SPRIselect Beads (Beckman-Coulter), and quality-checked with a D1000 ScreenTape assay on a TapeStation (Agilent) before sequencing. Libraries were pooled and sequenced on a NovaSeq X system at UCSF’s Center for Advanced Technology (CAT). For the IFN-γ screen, half of the tumor-isolated T cells were stimulated with CST Dynabeads at a 1.5:1 bead-to-cell ratio in the presence of Golgi Plug protein transport inhibitor (BD Biosciences, 555029, 1:1000 dilution). After 9 hours, cells were stained for surface markers, fixed, and processed for intracellular cytokine staining following BD Cytofix/Cytoperm kit instructions (BD Biosciences, 554714). The top 20% IFN-γ high and bottom 20% IFN-γ low populations were sorted at the Parnassus Flow Cytometry Core. Genomic DNA from these populations was extracted and processed as described above. After sequencing, FASTQ files were processed and analyzed using MAGeCK v0.5.9.5 as previously described^6^. For the abundance screen, sgRNA frequencies from spleen and tumor samples were compared to the input. For the IFN-γ screen, the IFN-γ high and low populations were compared. Guides with fewer than 40 reads in control samples were excluded from analysis.

### TIL isolation

For extraction of T cells from the tumors, the tumors were excised and digested in RPMI 1640 medium supplemented with Collagenase IV (1 mg/ml, Roche), DNase I (150 μg/ml, Roche), and 10% FBS. Digestion was carried out for 60 minutes at 37 °C with shaking at 200 rpm. The resulting digested tissues were then passed through 40μm cell strainers and subjected to further analysis. For isolation of TILs from ascites of ovarian cancer patients, we used the EasySep™ Human T Cell Isolation Kit (catalog no. 17951) following the manufacturer’s instructions, as described above.

### Pathway enrichment analysis

Pathway analysis was performed using the package clusterProfiler (v4.10.1) in R (v 4.3.3). The GSEA function was run using the msigdbr (v7.5.1) package as the source of the Homos sapiens Molecular Signatures Database. Pathway visualization was performed using clusterprofiler, enrichplot (v1.22.0), and ggplot2 (v3.5.1) in R with filtering for specific Gene Ontology (GO) categories. Manual annotation of GPCR pathway members guided by theusing literature was performed in addition to the canonical (GO) pathway annotation for rank plot visualization.

### In vitro Incucyte killing assay

IncuCyte live-cell imaging assays were performed using primary human CAR T cells, ovarian cancer patient-derived CAR TILs, and TILs isolated from tumor-bearing mouse models. T cells were co-cultured with pre-plated mKate⁺ CD19-expressing A375 cells, mKate⁺ Skov3 cells, or mKate⁺ A375 cells engineered to express an anti-CD3 scFv, in 384-well flat-bottom plates (Corning). T cells were seeded at various effector-to-target (E:T) ratios. Plates were imaged every 6 hours using the IncuCyte S3 Live-Cell Analysis System (Essen BioScience). Red fluorescent object counts (mKate⁺ cells) were recorded over time. Cancer cell growth was assessed for each replicate at each time point by normalizing red object counts to the initial time point (time zero).

### Multiplex immunoassay

To assess cytokine production in CAR T cells treated with GPCR agonists, CD19 CAR T cells were analyzed five days after CRISPR editing. Cells were resuspended in fresh cytokine-free cX-VIVO medium and co-cultured with CD19-expressing A375 cells at a 1:1 ratio for 24 hours under the GPCR agonist treatment conditions described above. Following co-culture, supernatants were collected, centrifuged at 500 × *g* for 5 minutes at 4 °C to remove cellular debris, aliquoted, and stored at −80 °C. Cytokine concentrations were quantified using the LEGENDplex™ Human CD8/NK Panel (13-plex) with VbP V02 (BioLegend, 741187) according to the manufacturer’s instructions. Data were analyzed using LEGENDplex™ analysis software (BioLegend).

### T cell proliferation and viability assays

To assess proliferation and viability, CRISPR-edited T cells were analyzed three days after removal of anti-CD3/CD28 Dynabeads. Cells were resuspended in fresh cX-VIVO medium and seeded at a density of 1 × 10⁶ cells/mL in 200 μL volumes into flat-bottom 96-well plates, with or without IL-2 supplementation. Cell counts and viability were measured using the Cellaca MX Automated Cell Counter (Nexcelom). At each passage, half of the culture volume was discarded and replaced with fresh medium, and the number of cells maintained was used to calculate total cell counts. For evaluation of proliferation and viability under GPCR agonist treatment, the indicated agonists were added at the concentrations described above.

### In vivo mouse tumor models

All mouse experiments were conducted in accordance with protocols approved by the Institutional Animal Care and Use Committee (IACUC) at UCSF.

For the in vivo human T cell studies, NOD.Cg-Prkdc^scid^ Il2rg^tm1Wjl^/SzJ (NSG, Strain #005557) mice were either purchased from The Jackson Laboratory (JAX) or bred in-house under sterile conditions at the UCSF Laboratory Animal Resource Center. Unless otherwise specified, 1 × 10⁶ tumor cells suspended in 100 μL of serum-free RPMI-1640 medium were injected subcutaneously into each mouse. Nine days post-engraftment, mice with comparable tumor burdens were randomized to receive intravenous (i.v.) infusion of 1 × 10⁶ CAR T or TCR T cells. For the B16 melanoma tumor model, female C57BL/6 mice aged 6–8 weeks purchased from JAX were injected subcutaneously with 3 × 10⁵ B16-OVA melanoma cells. Mice with similarly sized tumors were randomized to receive treatments of CARISPR Cas9-edited OT-I T cells. On days 10 post-tumor inoculation, 7 × 10⁶ of T cells was administered via i.v. Subcutaneous tumors were measured using digital calipers twice weekly for fast-growing tumor models and once weekly for slower-growing models. Tumor volume was calculated using the following formula: Tumor volume = (π × Length × Width × (Length + Width)) / 12. For subcutaneous tumor models, endpoint was defined as a progressively growing tumor reaching 20 mm in any dimension.

### Flow cytometry

For in vivo tumor models, cells were isolated from the spleen and tumor tissues. Erythrocytes were removed using 1X RBC lysis buffer (eBioscience), and dead cells were excluded using Fixable Viability Dye eFluor™ 780 (eBioscience). For surface marker analysis, cells were stained with specific antibodies in FACS buffer (1× PBS supplemented with 2% FBS and 2 mM EDTA) for 30 minutes at 4 °C. To assess cytokine expression in Figure 1 and Extended Data Figure 2, cells were stimulated with anti-CD3/CD28 Dynabeads at a 1:1 bead-to-cell ratio in the presence of Golgi Plug protein transport inhibitor for 6 hours. For IFN-γ expression analysis in the screen, tumor-infiltrating cells were stimulated with anti-CD3/CD28 Dynabeads at a 1.5:1 bead-to-cell ratio with Golgi Plug for 9 hours. After stimulation, cells were stained for surface markers and viability, then fixed and permeabilized using the BD Cytofix/Cytoperm™ kit, followed by intracellular cytokine staining. For analysis of cell surface activation markers, 200,000 T cells were seeded per well in a round-bottom 96-well plate and stimulated with ImmunoCult™ at 12.5 μL/mL for 6 hours at 37 °C. To assess mitochondrial membrane potential, cells were stained with 5 μM MitoTracker™ Red CMXRos (Invitrogen) in 1× HBSS for 30 minutes at 37 °C in 5% CO₂. Samples were acquired using either an Attune™ NxT Flow Cytometer (Invitrogen) or a BD LSRFortessa™ X-20 Cell Analyzer, and data were analyzed using FlowJo software. Antibodies used: Pacific Blue™ anti-human TNF-α (Biolegend, 502920), APC anti-human CD279 (Biolegend, 621610), Brilliant Violet 421™ anti-human CD366 (Biolegend, 345008), PE anti-human CD25 (Biolegend, 302606), Brilliant Violet 421™ anti-human CD69 (Biolegend, 310930), Brilliant Violet 711™ anti-human CD45 (Biolegend, 304050), PE anti-human CD62L (Biolegend, 304806), APC anti-human CD62L (Biolegend, 304810), Brilliant Violet 421™ anti-human CD45RA (Biolegend, 304130), PE anti-human CD45RA (Biolegend, 304108), APC anti-human TCR α/β (Biolegend, 306718), APC anti-human IFN-γ (Biolegend, 502512), PE mouse anti-Human IFN-γ (BD Pharmingen, 554701), FITC anti-human CD4 (Biolegend, 317408), APC anti-human CD4 (Biolegend, 317416), PE anti-human CD8a (Biolegend, 300908), APC anti-human CD137 (Biolegend, 309810), Brilliant Violet 711™ anti-human CD3 (Biolegend, 317328), PE anti-human CD39 (Biolegend, 328208), APC anti-DYKDDDDK Tag (Biolegend, 637308), PE anti-DYKDDDDK Tag (Biolegend, 637310), APC anti-human TCR Vβ13.1 (Biolegend, 362408), PE/Cyanine7 anti-human Ki-67 (Biolegend, 350526), Whitlow/218 Linker (E3U7Q) Rabbit mAb (Alexa Fluor® 488 Conjugate) (Cell signaling technology inc, 55809S).

## CRISPR knockout using electroporation

Human primary T cells were isolated, activated, and transduced with lentiviral CAR constructs as previously described. 48 hours post-activation, cells were counted and resuspended in Amaxa P3 Primary Cell solution (Lonza, V4SP-3960) at a concentration of 1–2 × 10⁶ cells per 20 µL. For CRISPR editing, cells were electroporated with 3.5 µL of ribonucleoprotein (RNP) complex using the Lonza 4D-Nucleofector with program EH115. Electroporated cells were immediately transferred into pre-warmed, cytokine-free human T cell medium and recovered at 37 °C for 30 minutes prior to transfer into complete culture medium. Anti-CD3/CD28 Dynabeads were removed two days after activation.

RNP complexes were generated using lyophilized crRNAs and tracrRNAs (Dharmacon), each resuspended in IDT duplex buffer (Cat# 1072570) to 160 µM. Equal molar amounts of crRNA and tracrRNA were annealed by incubation at 37 °C for 30 minutes. Cas9 protein (MacroLab; 40 µM stock) was added at a 1:2 molar ratio (Cas9: sgRNA duplex), along with 1 µL/well poly glutamic acid (PGA, 100 mg/mL stock), and the complex was incubated at 37 °C for 15 minutes before use. crRNA sequences used are listed in Extended Data Table 3.

To generate TRAC-targeted knock-in B7H3 CAR T cells, anti-CD3/CD28 beads were removed 48 hours post-activation. Cells were electroporated with RNP as described above, using sgRNA targeting the TRAC locus (Extended Data Table 3). 30 minutes post-electroporation, concentrated AAV particles encoding the B7H3 CAR construct were added to the cells, followed by overnight incubation in serum-free T cell medium. The following day, cells were resuspended in complete T cell culture medium and expanded initially at 2 × 10⁶ cells/mL, then maintained at 1.5 × 10⁶ cells/ml. Editing efficiency, CAR knock-in, and lentiviral transduction were assessed by flow cytometry prior to in vivo administration.

### Seahorse assays

Extracellular acidification rate (ECAR) and oxygen consumption rate (OCR) were measured using an XF96 Extracellular Flux Analyzer (Agilent) following the manufacturer’s protocols. Briefly, human T cells isolated from tumor-bearing NSG mice or treated with various GPCR agonists (as described above) were stimulated with ImmunoCult™ (Stemcell Technologies, 10971) at 12.5 µL/mL for 24 hours. A total of 200,000 cells per well were seeded onto XF96 microplates pre-coated with poly-D-lysine (Sigma, 103729-100). ECAR was measured using the Glycolysis Stress Test Kit (Agilent, 103020). After three baseline measurements, sequential injections of 11 mM glucose, 1.5 µM oligomycin, and 100 mM 2-deoxyglucose were performed. OCR was assessed using the Mito Stress Test Kit (Agilent, 102601). Following three baseline measurements, 2 µM oligomycin, 2 µM FCCP, and 0.5 µM rotenone/antimycin A were sequentially injected.

### Migration inhibition transwell bioassay

Transwell migration assays were performed as previously described using WEHI-231 cells, which are mixed with wild-type and P2RY8-GFP-expressing WEHI-231 cells^34^. Briefly, WEHI-231 cells were washed twice and resuspended at 2 × 10⁶ cells/mL with pre-warmed migration medium (RPMI 1640 supplemented with 0.5% fatty acid–free BSA, 50 IU/mL penicillin/streptomycin, and 10 mM HEPES), then incubated at 37 °C for 10 minutes. Recombinant human CXCL12 (PeproTech) was diluted to 50 ng/mL in the migration medium. Tumor cell supernatants were diluted at the indicated ratios (**Fig. 3d**) in CXCL12-containing migration medium, and 600 µL of each mixture was added to the lower chamber of a 24-well plate. Transwell inserts (6.5 mm diameter, 5 µm pore size; Corning) were placed into each well, and 100 µL of WEHI-231 cells (2 × 10⁵ cells) was added to the upper chamber. After a 3-hour incubation at 37 °C, cells in the lower chamber were collected and analyzed by flow cytometry.

To generate tumor cell supernatants, 1.5 × 10⁵ tumour cells were seeded per well in 6-well plates. Once confluent, the medium was replaced with 1.5 mL per well of migration medium. After 24 h, supernatants were collected and centrifuged to remove cellular debris.

### Migration assays of human T cells

For the in vitro transwell migration assay, AAVS1 control and P2RY8 KO T cells were labeled with CellTrace Violet and CellTrace^TM^ Far Red (Thermo Fisher Scientific) at a 1:2000 dilution, respectively, and mixed at a 1:1 ratio three days post-editing. The cells were resuspended in migration medium at approximately 2 ×10⁶ cells/mL and incubated at 37°C for 10 minutes. Recombinant human CXCL12 (Peprotech) was diluted to 50 ng/mL in tumor supernatant, and 600 µL of this mixture was added to the lower chamber of a 24-well plate. A total of 2 × 10⁵ T cells in 100 µL were added to the upper chamber of a 6.5-mm transwell insert (5 µm pore size; Corning). After a 3-hour incubation at 37°C, cells that migrated to the lower chamber were collected and quantified by flow cytometry.

For the in vivo migration assay, human primary T cells transduced with GFP-labeled control sgRNA or BFP-labeled sgRNA targeting P2RY8 were mixed at a 1:1 ratio and co-transferred into tumor-bearing NSG mice three weeks after tumor implantation. Forty-eight hours post-transfer, the relative proportions of GFP⁺ and BFP⁺ CAR T or TCR T cells were analyzed in the spleen and tumor by flow cytometry.

### GGG extraction and measurement

GGG levels in tumor cell culture supernatants were quantified by LC–MS/MS as previously described^34^. Briefly, after removing cells and debris, 5 mL of each tumor cell supernatant was acidified to pH 3, mixed with methanol to achieve a final concentration of 20% MeOH, and spiked with LTC4-d5 as an internal standard. Samples were loaded onto a 500-mg C18 solid-phase extraction (SPE) column (Agilent Technologies), washed twice with Milli-Q H₂O, eluted with methanol, dried, and reconstituted in 200 µL of methanol for analysis.

Samples were analyzed using a Shimadzu 30-AD UPLC in series with a SCIEX 7500 Triple Quadrupole Mass Spectrometer. A 5 µL aliquot was loaded onto a Synergi Polar-RP column (75 × 4.6 mm) kept at 40 °C, with a mobile phase gradient of A: H₂O + 0.1% formic acid and B: acetonitrile + 0.1% formic acid. The gradient was programmed as follows: 0-1 min, 40% B; 1-4 min, ramp to 95% B; 4-6 min, hold at 95% B; 6–6.5 min, ramp to 50% B; 6.5-8 min, return to 40% B. GGG was detected in positive ionization mode using multiple reaction monitoring (MRM) with a transition of m/z 580.3 → 179.0. The internal standard LTC4-d5 was detected using the transition m/z 631.4 → 179.0.

Instrument settings were as follows: source temperature 400 °C, curtain gas (CUR) 40, ion source gas 1 (GS1) 40, gas 2 (GS2) 70, ion spray voltage (IS) 2000 V. Compound-specific settings were: Entrance potential (EP) 10 V, collision cell exit potential (CXP) 10 V, collision energy (CE) 35 eV, dwell time 100 ms, and pause time 5.007 ms. Peak area was integrated using Sciex OS software (version 2.1.6.59781).

## Statistical Information

Statistical analysis was performed using Prism 10 software (version 10.2.3). Results are presented as mean ± s.e.m. A *P* value ≤ 0.05 was considered statistically significant. Unless otherwise specified, a two-tailed unpaired Student’s *t*-test was used for comparisons between two groups, two-way ANOVA was used for tumor growth curve analysis, and the log-rank (Mantel–Cox) test was applied for survival comparisons.

## Acknowledgements

We thank members of the Carnevale, Marson, and Eyquem laboratories, as well as T. Tolpa, for their valuable support. We are also grateful to the UCSF Flow Cytometry Core and the Mouse Metabolism Core (supported by NORC grant P30DK098722) for their technical assistance. J.C. received funding from the Parker Institute for Cancer Immunotherapy, CRISPR Cures for Cancer, the Pascarella Scholars fund, the Burroughs Wellcome Fund and the Lydia Preisler Shorenstein Donor Advised Fund. Q.L. was supported by the Outstanding Doctoral Graduates Development Scholarship of Shanghai Jiao Tong University. A.M. received funding from the Simons Foundation, the Lloyd J. Old STAR Award (Cancer Research Institute), the Parker Institute for Cancer Immunotherapy, the Innovative Genomics Institute, the Larry L. Hillblom Foundation (grant 2020-D-002-NET), the Arc Institute, the Byers family, K. Jordan, and the CRISPR Cures for Cancer Initiative.

## Author Contributions

Q.L., J.C, and A.M. conceived and designed the study. Q.L. developed the in vivo screening platform, performed the CRISPR screens, primary T cell in vitro and in vivo experiments, and analyzed the data. P.A.C., E.U., S.M.Z., and T.N.L. assisted with in vitro CAR T validation and migration assays. P.A.C., E.U., S.M.Z., and Z.M.L. contributed to in vivo CAR T experiments, N.K. made the Brunello library virus and F.L.P. assisted with the in vivo migration assay. C.H.W. conducted the P2RY8 evolutionary analysis. G.M.A. and K.F. provided the B7H3 CAR construct and ascites samples from ovarian cancer patients, respectively. Q.L., M.M.A. and E.S. performed the computational analyses. Q.L. prepared the figures with input from J.C., S.E.D., and A.M. J.C., S.E.D., and Q.L. wrote the manuscript, with input and editing from all authors.

## Additional Information

Supplementary Information is available for this paper. Correspondence and requests for materials should be addressed to Qi Liu or Julia Carnevale.

## Data Availability

All CRISPR screen data generated in this study are provided in Extended Data Tables 1 and 2. The crRNA sequences used are listed in Extended Data Table 3. Source data are available with this paper.

## Tables

Table 1. MAGeCK analysis results from in vivo genome-wide CRISPR knockout abundance screens

Table 2. MAGeCK analysis results from in vivo genome-wide CRISPR knockout IFN-γ screens

Table 3. crRNA sequences used in this study

Table 4. Information on primary human T cell donors used in in vivo screening and validation studies

---

**Extended Data Fig. 1.**
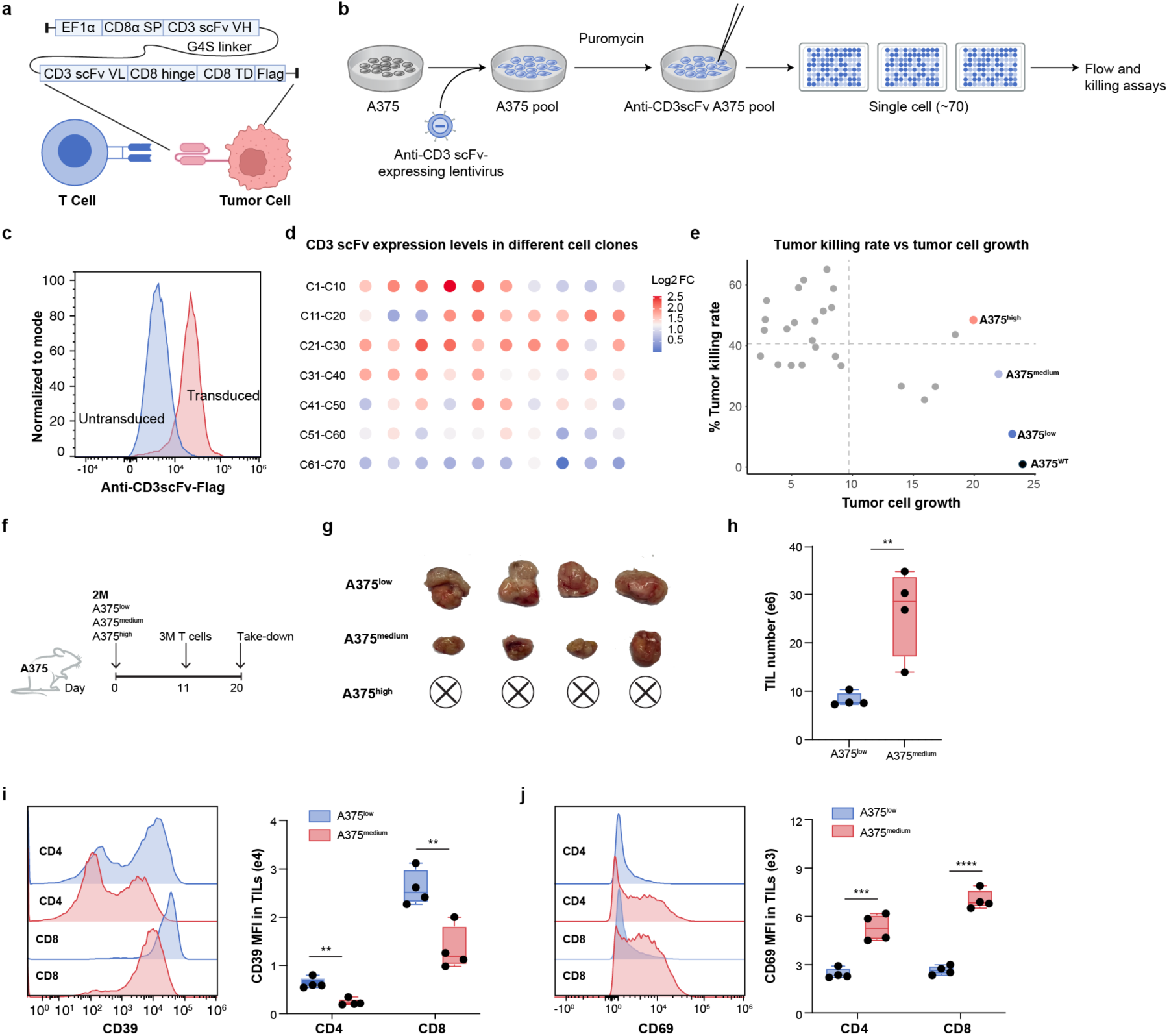
Clonal selection and evaluation to establish a model system with high T cell recovery from tumors for screening. **a.** Schematic of the in vivo model using tumor cells engineered to express an anti-CD3 scFv for intratumoral enrichment of human T cells. Also shown are the components of the lentiviral construct used to express the anti-CD3 scFv. **b.** Workflow depicting the generation of anti-CD3 scFv overexpressing A375 single cell clones. **c.** Flow cytometric analysis of anti-CD3 scFv surface expression on bulk-transduced vs untransduced A375 cells. **d.** Anti-CD3 scFv expression level as assessed by flow cytometry in 70 selected single cell clones. **e.** Scatter plot showing Incucyte-based tumor cell killing rates of A375 cell lines in the presence of primary human T cells plotted vs. rate of tumor growth of each line in the absence of T cells. Tumor lines include wild-type A375 cells and 27 of the anti-CD3 scFv-overexpressing A375 single-cell clones selected from (d). For clones with comparable anti-CD3 scFv expression levels, a single representative clone was selected for further analysis. Labeled clones were assessed in follow-up experiments. **f-j.** A375^low^, A375^medium^, and A375^high^ single-cell clones from (e) were subcutaneously engrafted into NSG mice, followed by intravenous infusion of 3M primary human T cells. Shown are: the experimental timeline (**f**), visual of tumor size at day 20 post-engraftment (**g**), absolute number of tumor-infiltrating lymphocytes (TILs) at day 20 (**h**), and expression of CD39 and CD69 in CD4⁺ and CD8⁺ TILs (**i-j**, n = 4 mice per group). Note that the A375^high^ tumors were fully eradicated, and thus we were unable to phenotype any T cells from those tumors in **h-i**. *P* values were determined by two-tailed unpaired Student’s t-test (**h, i, j**). \*\**P* < 0.01, and \*\*\*\**P* < 0.0001. Data are presented as mean ± s.e.m.

**Extended Data Fig. 2.**
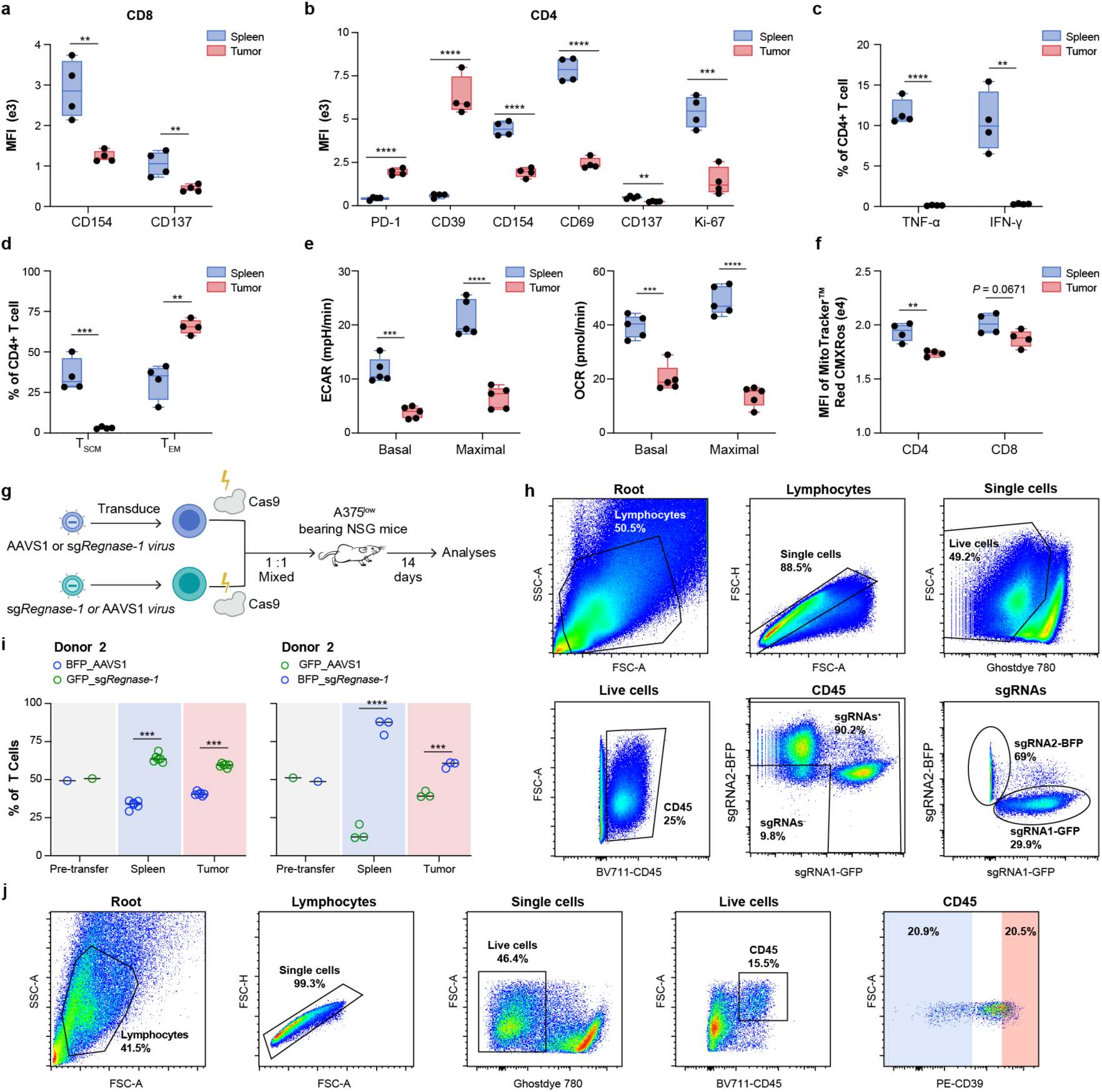
Phenotypic analysis of T cells from CD3 scFv-expressing tumors compared to spleens. **a-f.** The A375^low^ single-cell clone was subcutaneously engrafted into NSG mice, followed by intravenous infusion of primary human T cells. Comparative analyses of T cell phenotypes in spleen versus tumor were performed 9 days post-treatment. Shown are: MFI of CD154 and CD137 in CD8⁺ T cells (**a**); MFI of PD-1, CD39, CD154, CD69, CD137, and Ki-67 in CD4⁺ T cells (**b**); percentage of TNF-α⁺ and IFN-γ⁺ CD4⁺ T cells (**c**); percentage of Tscm and Tem CD4⁺ T cells (**d**); statistical analysis of ECAR and OCR (**e**), and MFI of MitoTracker™ Red CMXRos staining (**f**) in T cells isolated from spleen and tumor (n = 4 mice for **a-d** and **f**; n = 5 technical replicates for **e**, representative data from one of three mice for **e**). Note that a full complement of assays was completed on all CD4 and CD8 T cells, but these data are spread between Fig. 1 and here in Extended Data Fig. 2. **g-i.** In vivo competition assay of AAVS1 control and Regnase-1 KO primary human T cells in A375^low^ tumor-bearing mice. Shown are: the experimental timeline (**g**); flow cytometry gating strategy **(h)** and statistical analysis of the relative abundance of AAVS1 control vs Regnase-1 sgRNA-transduced T cells among recovered sgRNA⁺ CD45⁺ T cells from a second donor (**i**). Left: BFP_AAVS1 control sgRNA mixed with GFP_Regnase-1 sgRNA; Right: GFP_AAVS1 control sgRNA mixed with BFP-Regnase_1 sgRNA (n = 3-4 mice for each group). **j.** Flow cytometry gating strategy for sorting TILs based on CD39 expression: top (CD39^+^) and bottom (CD39^-^) 20% of TILs were sorted from A375^low^ tumor-bearing NSG mice. *P* values were determined by two-tailed unpaired Student’s t-test (**a-f, i**). \*\**P* < 0.01, \*\*\**P* < 0.001, and ****0.0001. Data are presented as mean ± s.e.m.

**Extended Data Fig. 3.**
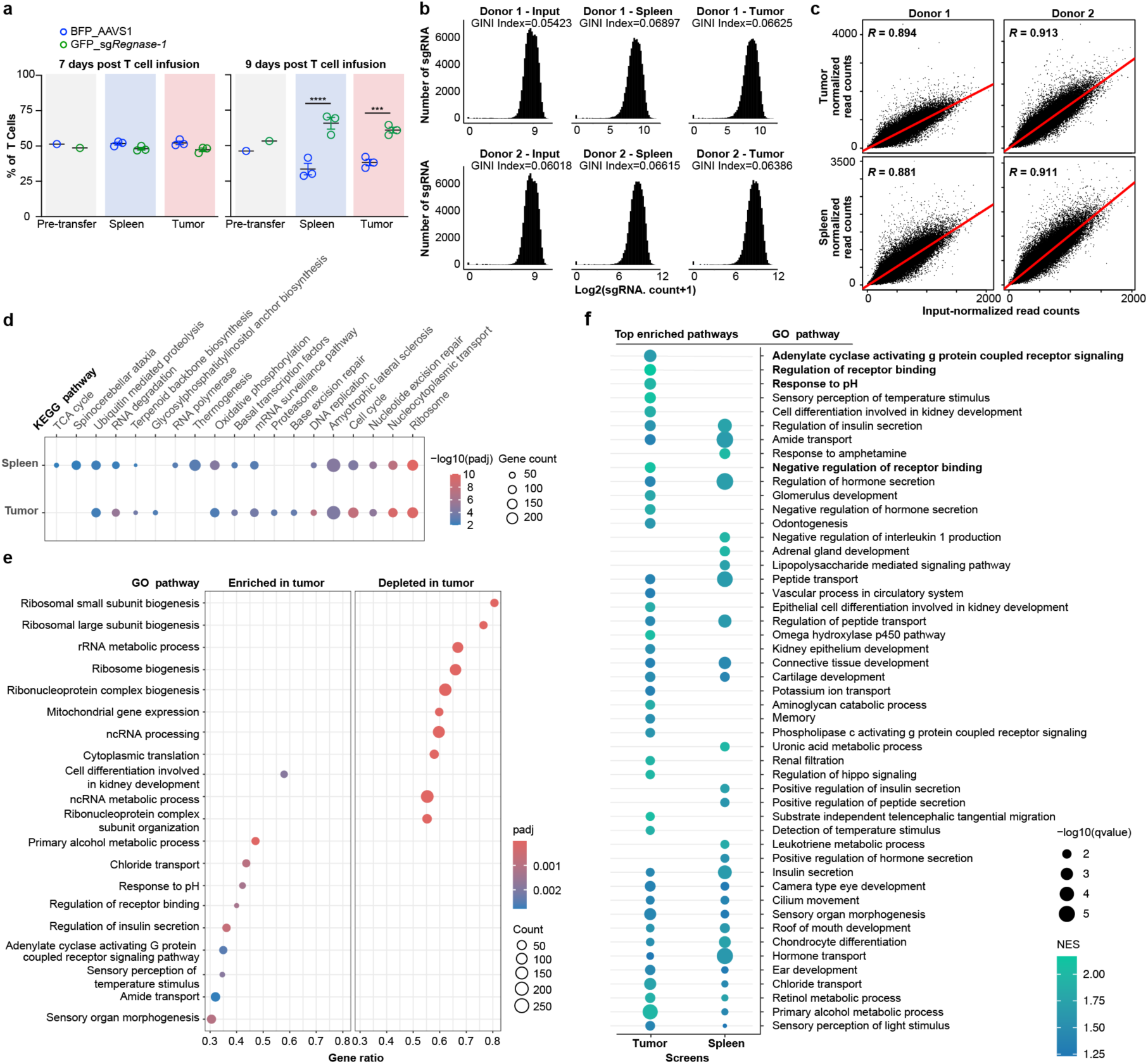
Quality control metrics and pathway enrichment analysis for the genome-wide in vivo CRISPR screen of T cell abundance. a. In vivo competition assay of AAVS1 control and Regnase-1 KO primary human T cells. T cells were analyzed 7 days (left) and 9 days (right) post-infusion (n = 3 mice per group). b. Distribution and GINI Index of sgRNA read counts from deep sequencing of T cells isolated from input, spleen, and tumor samples across two donors in the *in vivo* abundance screens. c. Pearson correlation of normalized sgRNA read counts comparing tumor versus input and spleen versus input across two donors in the in vivo abundance screen. d. KEGG pathway enrichment analysis of dropout genes in the tumor and spleen abundance screens. The top 20 enriched KEGG pathways (ranked by adjusted p-value) were selected independently for each screen. Pathways associated with diseases (e.g., cancer, infection, neurodegeneration) were removed to focus on canonical signaling and metabolic pathways. Dot size indicates the number of genes involved; color reflects statistical significance (-log_10_ adjusted *p*-value). e. Gene Ontology (GO) pathway analysis showing enriched pathways among the top enriched and depleted gene hits from the tumor abundance screen (tumor versus input). Points are scaled to the number of gene counts in different pathways and padj represents adjusted *P*-value. f. GO biological process analysis showing enriched pathways in the tumor (tumor vs. input) and spleen (spleen vs. input) abundance screens. Enrichment was based on NES with a minimum q-value < 0.0005. Dot size represents-log_10_(q-value). *P* values were determined by two-tailed unpaired Student’s t-test (**a**). \*\*\**P* < 0.001, and \*\*\*\**P* < 0.0001. Data are presented as mean ± s.e.m.

**Extended Data Fig. 4.**
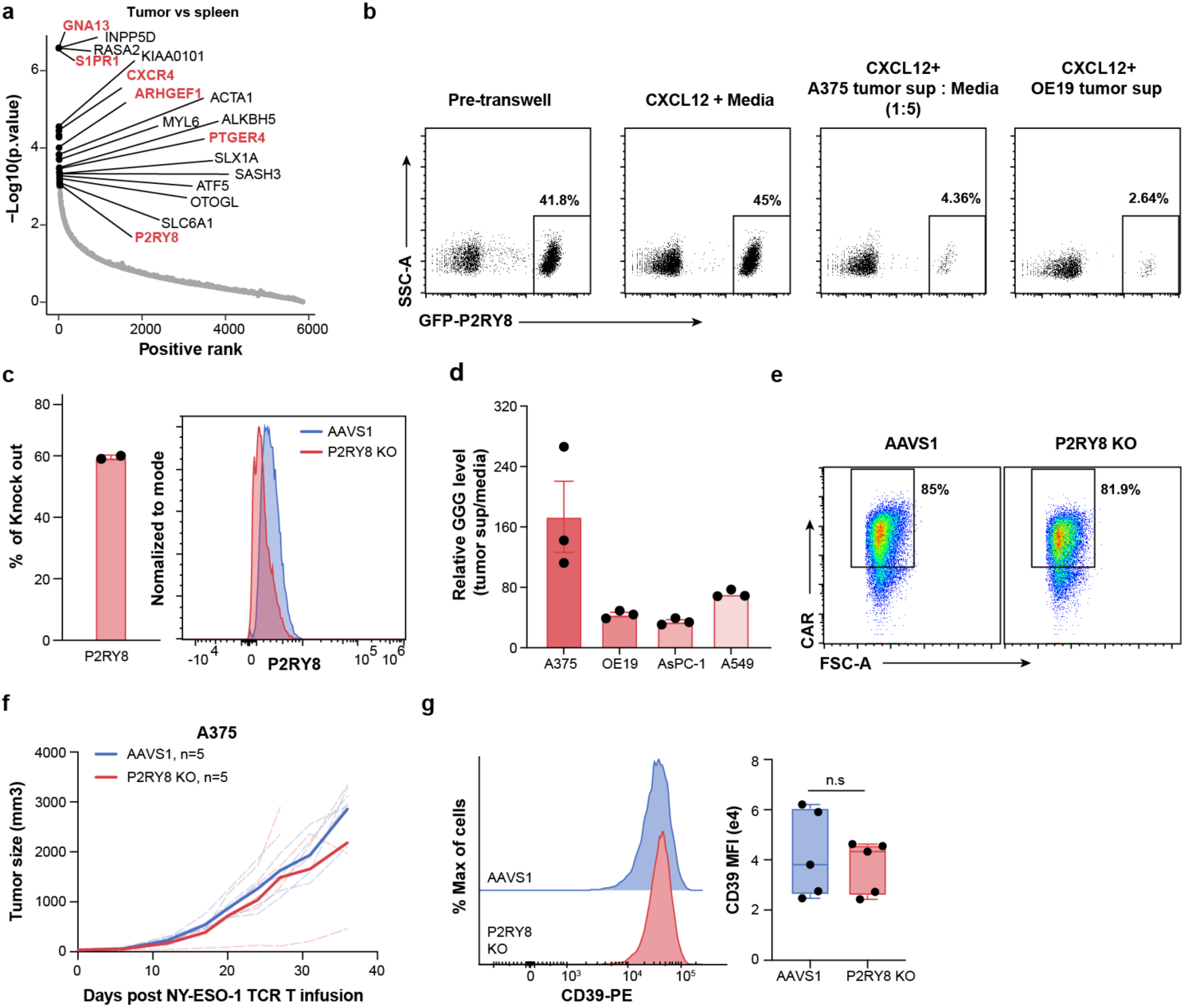
P2RY8 KO T cells have a competitive tumor infiltration advantage. **a.** Enriched gene ranks in tumor versus spleen across two primary human T cell donors, as determined by MAGeCK analysis. Genes depleted in the tumor versus input comparison were excluded. The top 30 ranked genes are highlighted as black dots. Genes with known roles in GPCR signaling or previously characterized functions in cell trafficking and migration are labeled in red. **b.** Representative flow cytometry plots from the P2RY8 ligand bioassay using 50 ng/mL CXCL12 and a mixture of WT and P2RY8-GFP⁺ WEHI-231 cells, measuring the percentage of cells that migrated to the bottom well, corresponding to Fig. 3d. **c.** CRISPR editing efficiency of the P2RY8 gene in primary human T cells, as determined by knockout score from ICE analysis (left) and P2RY8 protein expression measured by flow cytometry (right). **d.** Relative GGG levels in the supernatants of the indicated tumor cell lines compared to media control, as assessed by mass spectrometry analysis (n = 3 biological replicates). **e.** Flow cytometric histograms of surface CD19 CAR expression level in AAVS1 and P2RY8 KO CAR T cells prior to transfer to tumor-bearing NSG mice. **f-g.** A375 tumor-bearing NSG mice were treated with NY-ESO-1 TCR T cells 9 days post tumor injection. Shown are: (**f**) tumor growth kinetics, and (**g**) flow cytometric analysis of CD39 expression in tumor-infiltrating TCR^+^CD8⁺ T cells from AAVS1 control and P2RY8 knockout groups (n = 5 mice per group). *P* values were determined by two-tailed unpaired Student’s t-test (**g**). n.s, not significant. Data are presented as mean ± s.e.m.

**Extended Data Fig. 5.**
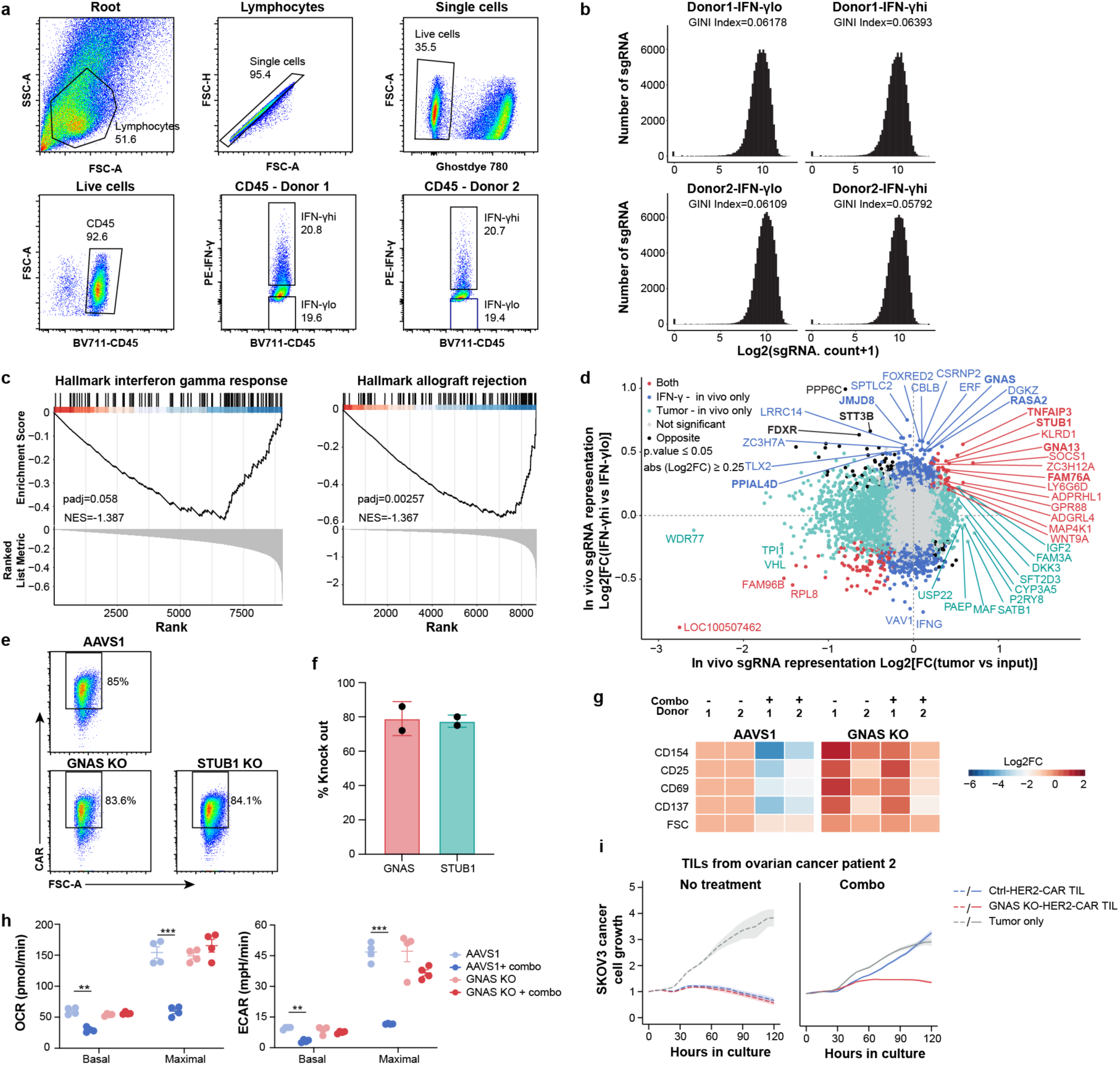
IFN-γ-based genome-wide in vivo CRISPR screen nominates genes regulating intratumoral T cell effector function. **a.** Flow cytometry gating strategy for sorting the top 20% IFN-γ high (IFN-γhi) and bottom 20% IFN-γ low (IFN-γlo) TILs from A375^low^ tumor-bearing NSG mice across 2 human T cell donors for the screen. TILs from the same donor were pooled for sorting. **b.** Distribution and GINI Index of sgRNA read counts from deep sequencing of TILs from IFN-γhi and IFN-γlo populations across two donors in the in vivo IFN-γ screens. **c.** GSEA of hallmark interferon gamma response and hallmark allograft rejection pathways in IFN-γhi versus IFN-γlo dropout genes. Data were analyzed across 2 human T cell donors. **d.** Comparison of in vivo primary human T cell screens, plotting Log2FC values from IFN-γ screen (IFN-γhi versus IFN-γlo) on the Y axis vs. Log2FC values from in vivo tumor abundance screen (tumor versus input) on the X axis. Data were analyzed across 2 human T cell donors. Genes shown in bold are those that also appear in the IFN-γ-in vivo only section of the comparison plot in Fig. 4c. P. value ≤ 0.05 and abs (Log2FC) ≥ 0.2. **e.** Flow cytometric histograms of surface CD19-CAR expression level in AAVS1, GNAS KO and STUB1 KO CD19 CAR T cells prior to transfer into tumor-bearing NSG mice. **f.** CRISPR editing efficiency of the GNAS and STUB1 genes in primary human T cells, as determined by knockout score from ICE analysis (n = 2 human T cell donors). **g-h.** AAVS1 and GNAS KO T cells were treated with a combination of the four agonists shown in Fig. 4g for 2 days, data shown are log_2_ fold change (day 2/day 0) of the MFI of indicated activation markers (**g**) and statistical analysis of OCR and ECAR (**h**, n = 2 human T cell donors for **g**, n = 4 technical replicates for **h**, and data are representative of one of three donors for **h**). **i.** Incucyte-based killing assay comparing GNAS KO and AAVS1 control HER2 CAR TILs in the presence (right) or absence (left) of the same four-agonist combination shown in Fig. 4g. TILs were isolated from the ascites of a second ovarian cancer (patient 2) and engineered to express HER2 CAR prior to co-culture with SKOV3 cancer cells. Parallel data for patient 1 is shown in Fig. 4g. *P* values were determined by two-tailed unpaired Student’s t-test in (**h**). \*\**P* < 0.01 and \*\*\**P* < 0.005. Data are presented as mean ± s.e.m.

**Extended Data Fig. 6.**
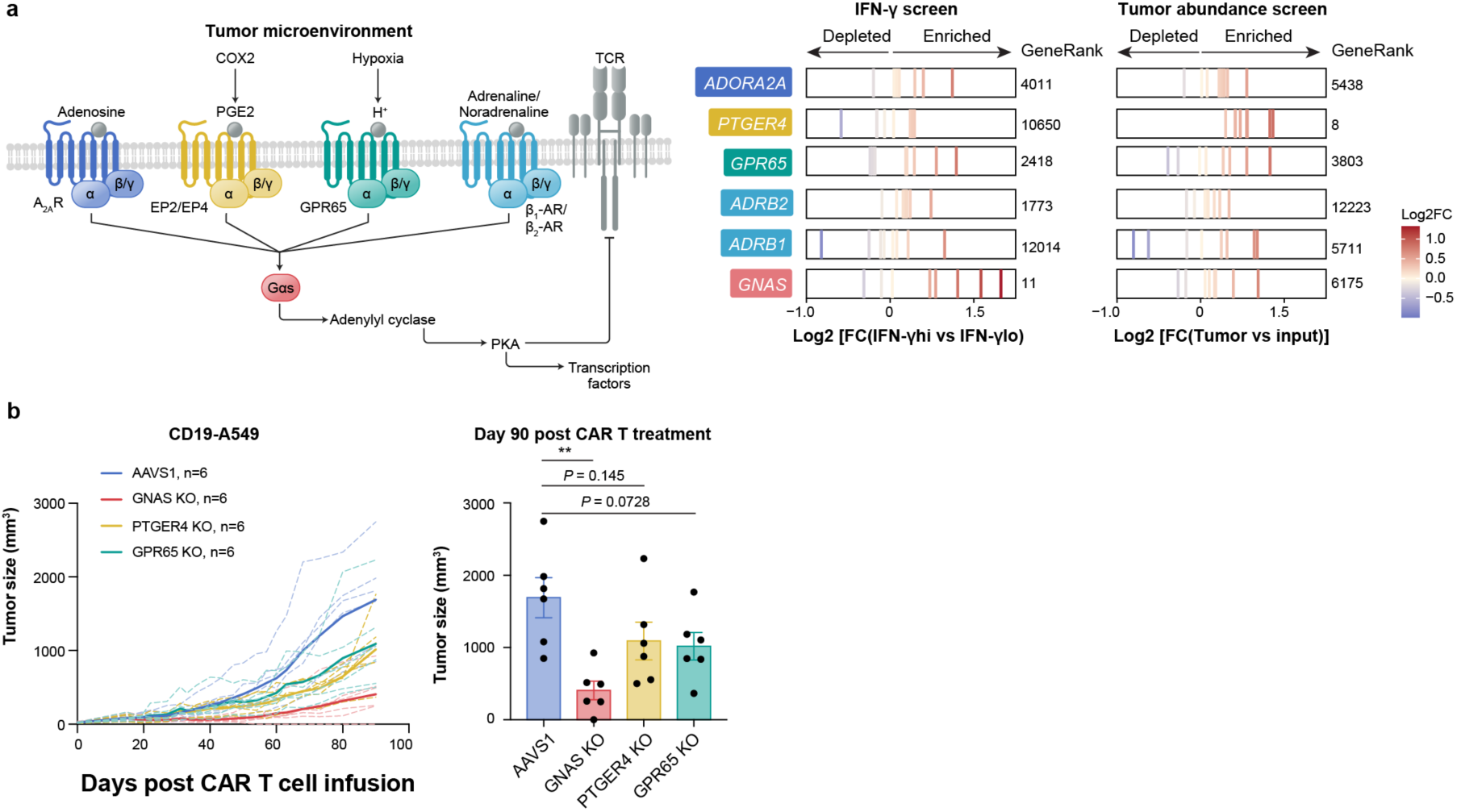
Analysis of GPCR genes upstream of Gαs across in vivo CRISPR screens and testing in preclinical models. a. sgRNA log_2_ fold changes for GPCR genes upstream of Gαs and gene rank values from the in vivo IFN-γ screen and tumor abundance screen. b. Tumor growth (left) and tumor size at day 90 post CAR T infusion (right) in CD19-A549 tumor-bearing NSG mice treated with AAVS1 control, GNAS KO, PTGER4 KO, or GPR65 KO CD19 CAR T cells, 9 days after tumor injection (n = 6 mice per group). *P* values were determined by two-tailed unpaired Student’s t-test in (**b**). \*\**P* < 0.01. Data are presented as mean ± s.e.m.

**Extended Data Fig. 7.**
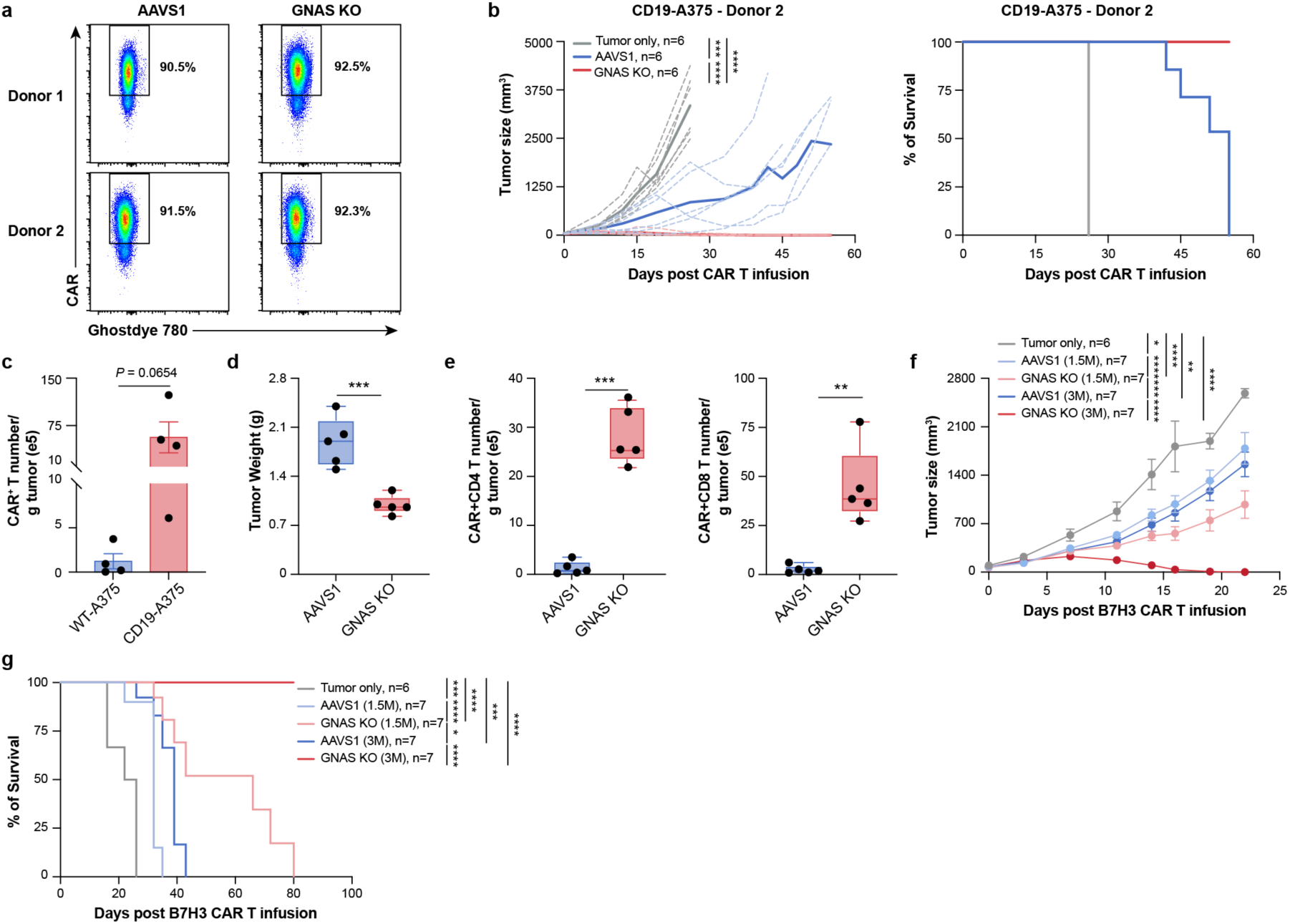
GNAS KO in therapeutic T cells enhances tumor control in multiple solid tumor preclinical models. **a.** Flow cytometric histograms of surface CD19 CAR expression level in AAVS1 and GNAS KO CD19 CAR T cells across 2 human T cells donors prior to transfer into tumor-bearing NSG mice. **b.** CD19-A375 melanoma cells were subcutaneously engrafted into NSG mice, followed by intravenous infusion of CD19 CAR T cells. Tumor growth curves (left), and mouse survival (right) are shown (n = 6 mice per group). **c.** Absolute number of CAR⁺ T cells per gram of WT-A375 or CD19-A375 tumors from the rechallenge experiment shown in Fig. 5b (n = 4 mice). **d-e.** Absolute tumor weight (**d**) and CAR^+^ CD4 and CD8 T cell number per gram of tumor (**e**) from the experiment shown in Fig. 5c (n = 5 mice each group). **f-g.** MES-SA uterine sarcoma cells were subcutaneously engrafted into NSG mice, followed by intravenous infusion of the indicated doses of AAVS1 or GNAS KO B7H3 CAR T cells. Tumor growth (**f**) and mouse survival (**g**) are shown (n = 6 mice for the tumor-only control group and n = 7 mice for all other groups). *P* values were determined by two-way ANOVA in (**b, f**) for tumor growth analysis, Log-rank (Mantel-Cox) test in (**b, g**) for mice survival analysis, and two-tailed unpaired Student’s t-test in (**c-e**). \**P* < 0.05, \*\**P* < 0.01, \*\*\**P* < 0.001, and \*\*\*\**P* < 0.0001. Data are presented as mean ± s.e.m.

**Extended Data Fig. 8.**
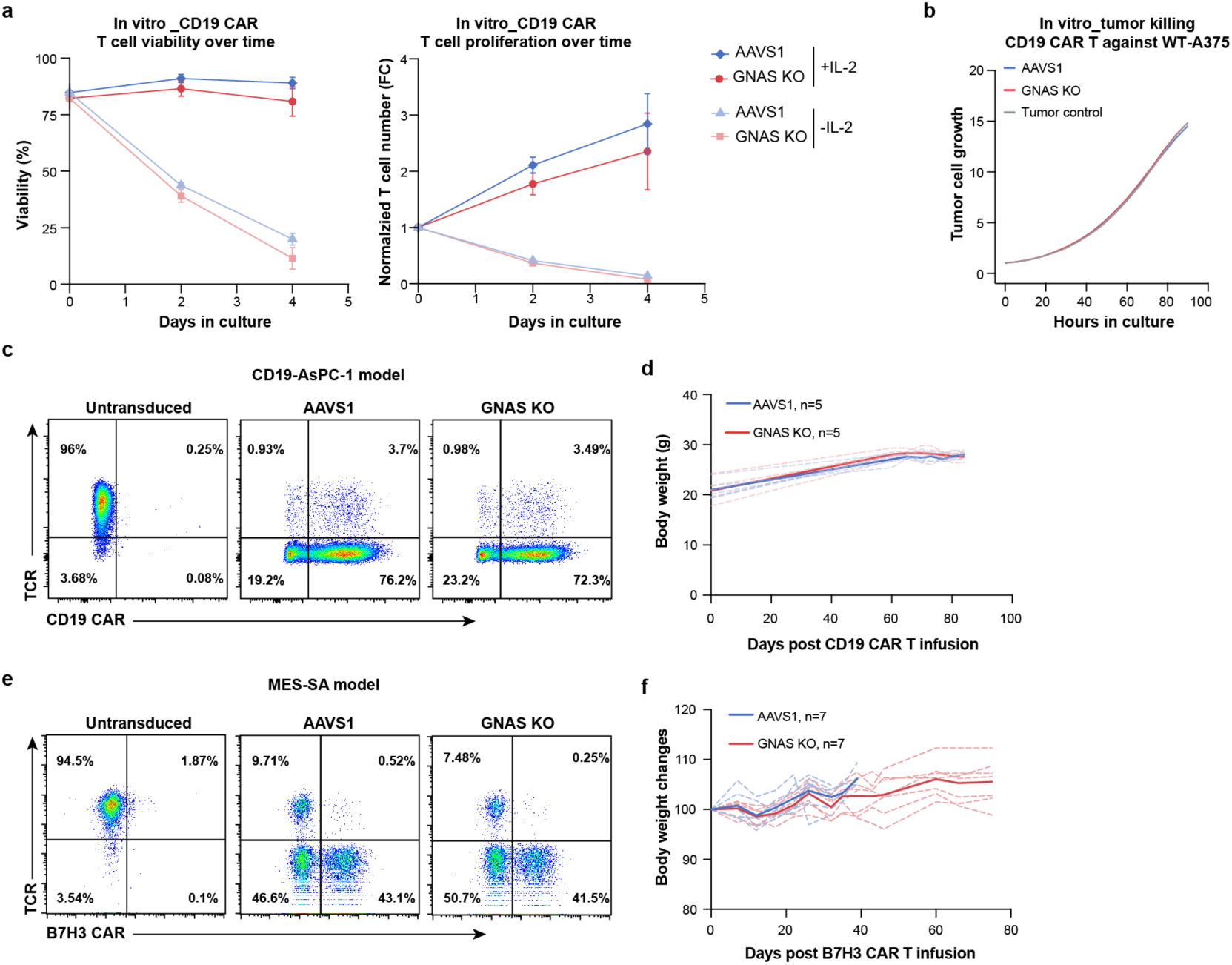
GNAS KO therapeutic T cells show no evidence of dysregulated proliferation, antigen-negative cell killing, or poor health indicators in mice. **a.** Cell viability (left) and normalized fold change in absolute T cell numbers (right) of CD19 CAR T cells cultured with or without cytokines over time. Fold change was normalized to T cell number on day 0 (n = 4 human T cell donors). **b.** Normalized tumor cell growth of wild-type A375 melanoma cells (antigen-negative) co-cultured with AAVS1 or GNAS KO CD19 CAR T cells at a 1:1 ratio, or cultured alone. Tumor growth was monitored using Incucyte live-cell imaging of mKate⁺ A375 cells. Data are representative of one of four donors. **c-d.** Flow cytometric histograms of surface expression of CD19 CAR and endogenous TCR in untransduced, AAVS1, and GNAS KO CAR T cells (**c**), and body weight of mice (**d**) from the experiment shown in Fig. 5f. **e-f.** Flow cytometric histograms of surface expression of B7H3 knock-in CAR and endogenous TCR in untransduced, AAVS1, and GNAS KO CAR T cells (**e**), and body weight of mice (**f**) from the experiment shown in Fig. 5g.

